# Neuroendocrine prostate cancer converges on a fetal pulmonary neuroendocrine-like program

**DOI:** 10.64898/2026.07.07.737089

**Authors:** Wenchang Yue, Natasha Kyprianou, Ashutosh K. Tewari, Babu J. Padanilam

**Affiliations:** Department of Urology, Icahn School of Medicine at Mount Sinai, New York, NY, USA; Tisch Cancer Institute, Icahn School of Medicine at Mount Sinai, New York, NY, USA; Department of Pathology and Molecular and Cell-Based Medicine, Icahn School of Medicine at Mount Sinai, New York, NY, USA; Department of Oncological Sciences, Icahn School of Medicine at Mount Sinai, New York, NY, USA

**Keywords:** Prostatic Neoplasms, Cell Plasticity, Cell Lineage, Fetal Development, Single-Cell Gene Expression Analysis, Gene Regulatory Networks, Spatial Transcriptomics

## Abstract

Advanced prostate cancer can relapse as neuroendocrine prostate cancer (NEPC), a treatment-resistant, androgen receptor (AR)-independent state whose developmental identity remains unclear. We integrated single-cell, bulk, cistrome and spatial datasets from multiple cohorts and mapped malignant states to human fetal atlases. In the discovery cohort (35,696 tumor cells, 20 patients), NEPC cells were malignant and shared copy-number architecture with adenocarcinoma (chromosome-level r = 0.64). NEPC most closely matched a fetal, lineage-restricted neuroendocrine program, best supported as pulmonary neuroendocrine-like among alternatives tested (islet, chromaffin, sympathoblast), with consistent direction across three cohorts; the pulmonary-versus-islet distinction was method-sensitive and remains provisional. Stage and pseudotime analyses supported an ordered luminal-to-neuroendocrine sequence, with FOXA2/SOX2 early and ASCL1/NEUROD1/MYCN late across five datasets. A regulatory screen nominated a circuit centered on ASCL1 whose convergent enhancers overlapped fetal pulmonary neuroendocrine chromatin and showed FOXA1 binding in cell-line and xenograft models (87.0% versus 33.8%; P = 0.016). In one NEPC Visium section, the program localized to malignant territory defined by copy number (AUROC 0.85). Based on three NEPC patients with cross-cohort replication, these data support NEPC as an adenocarcinoma-related malignant state converging on a fetal pulmonary neuroendocrine-like program and nominate ASCL1-linked regulatory and surface targets.

## Introduction

Most men with advanced prostate cancer respond initially to AR pathway inhibition, yet some relapse with tumors that have shed their luminal, AR-driven identity and acquired a neuroendocrine phenotype. NEPC is AR-independent, responds poorly to available therapy, and is associated with short survival [1,2]. Treatment-emergent small-cell neuroendocrine carcinoma is found in approximately 17% of men with metastatic castration-resistant prostate cancer treated with potent AR pathway inhibitors [2], and this phenotype has become increasingly clinically relevant in the current treatment era. Which neuroendocrine state an adenocarcinoma enters, and what transition accompanies that change, are therefore central questions for understanding AR pathway resistance.

Treatment-emergent NEPC is generally thought to arise from adenocarcinoma through lineage plasticity rather than from a separate cell of origin. NEPC and the adenocarcinoma it follows share genomic alterations consistent with a common clonal origin [1], and combined loss of *RB1* and *TP53* renders luminal prostate cells competent to transdifferentiate toward a neuroendocrine state, a switch supported by SOX2 and related factors [3,4]. The resulting state expresses canonical neuroendocrine regulators, including ASCL1, NEUROD1, INSM1 and MYCN, the last an established EZH2-linked driver of NEPC [5], with relief of the REST repressor, and ASCL1 is required for neuroendocrine differentiation in prostate cancer models [6]. These transcription factors are shared with normal neuroendocrine development and with small-cell lung cancer (SCLC), where programs driven by ASCL1, NEUROD1, POU2F3 and YAP1 define distinct subtypes [7,8].

Two connected aspects of this neuroendocrine program remain underdefined. First, it is usually described generically, as the presence of neuroendocrine markers; recent work has begun to resolve NEPC into ASCL1- and NEUROD1-defined subtypes [9,10], but which developmental neuroendocrine identity the tumor resembles has remained unspecified. Cancers can re-access developmental and fetal transcriptional programs as a route to plasticity and therapy resistance [11,12], and human fetal cell atlases now make it possible to ask, at single-cell resolution, which fetal lineage a tumor state matches [13,14]. For NEPC, this question has not been addressed by systematically integrating single-cell, chromatin and spatial evidence, and candidate lineages such as pulmonary neuroendocrine, pancreatic islet and neural crest derivatives remain undistinguished. Second, the order in which the luminal-to-neuroendocrine switch proceeds across patients, and the regulatory architecture associated with a program centered on ASCL1, are incompletely mapped, although FOXA1 and FOXA2 have been implicated as pioneer factors in the transition [15,16]. Whether the program can be localized in situ to malignant tissue, rather than inferred from dissociated cells, has also not been established.

Here we characterize the neuroendocrine identity of NEPC by integrating single-cell, bulk, cistrome and spatial data across multiple cohorts and mapping the malignant compartment against human fetal cell atlases [13,14]. NEPC cells were malignant and shared copy-number architecture with prostate adenocarcinoma in the discovery cohort, and their transcriptional state most resembled a fetal, lineage-restricted, pulmonary neuroendocrine-like program rather than a generic neuroendocrine one. The luminal-to-neuroendocrine switch was consistent with an ordered sequence across cohorts. An unbiased regulatory screen nominated a circuit centered on ASCL1 with FOXA1 binding at fetal pulmonary neuroendocrine chromatin, and spatial transcriptomics localized the program in situ to malignant NEPC territory. These findings refine NEPC from a generic neuroendocrine phenotype to a specific developmental reference state and nominate candidate dependencies and surface targets linked to ASCL1 for further study [6,17].

## Results

### NEPC malignant cells share copy-number architecture with adenocarcinoma and show a luminal-to-neuroendocrine shift

In the discovery cohort, 35,696 tumor cells from 20 patients (17 castration-resistant adenocarcinoma [CRPC], 3 NEPC) resolved into adenocarcinoma/CRPC (24,089 cells) and NEPC (11,607 cells) populations on the integrated Uniform Manifold Approximation and Projection (UMAP) (Fig. 1A); per-patient comparisons used the 13 patients with ≥100 cells (10 adenocarcinoma, 3 NEPC). Within the NEPC compartment, fetal atlas argmax labels were enriched for neuroendocrine identities (Fig. 1B). At the patient level, the three NEPC cases lost the luminal/AR program (*AR, KLK3, KLK2, NKX3-1, FOLH1*) and gained neuroendocrine markers (*CHGA, CHGB, SYP, NCAM1, INSM1, ASCL1, NEUROD1*), whereas adenocarcinoma patients kept it (Fig. 1C). Each patient contributed one aggregated value, so this was a per-patient, not cell-level, contrast.

**Figure 1.**
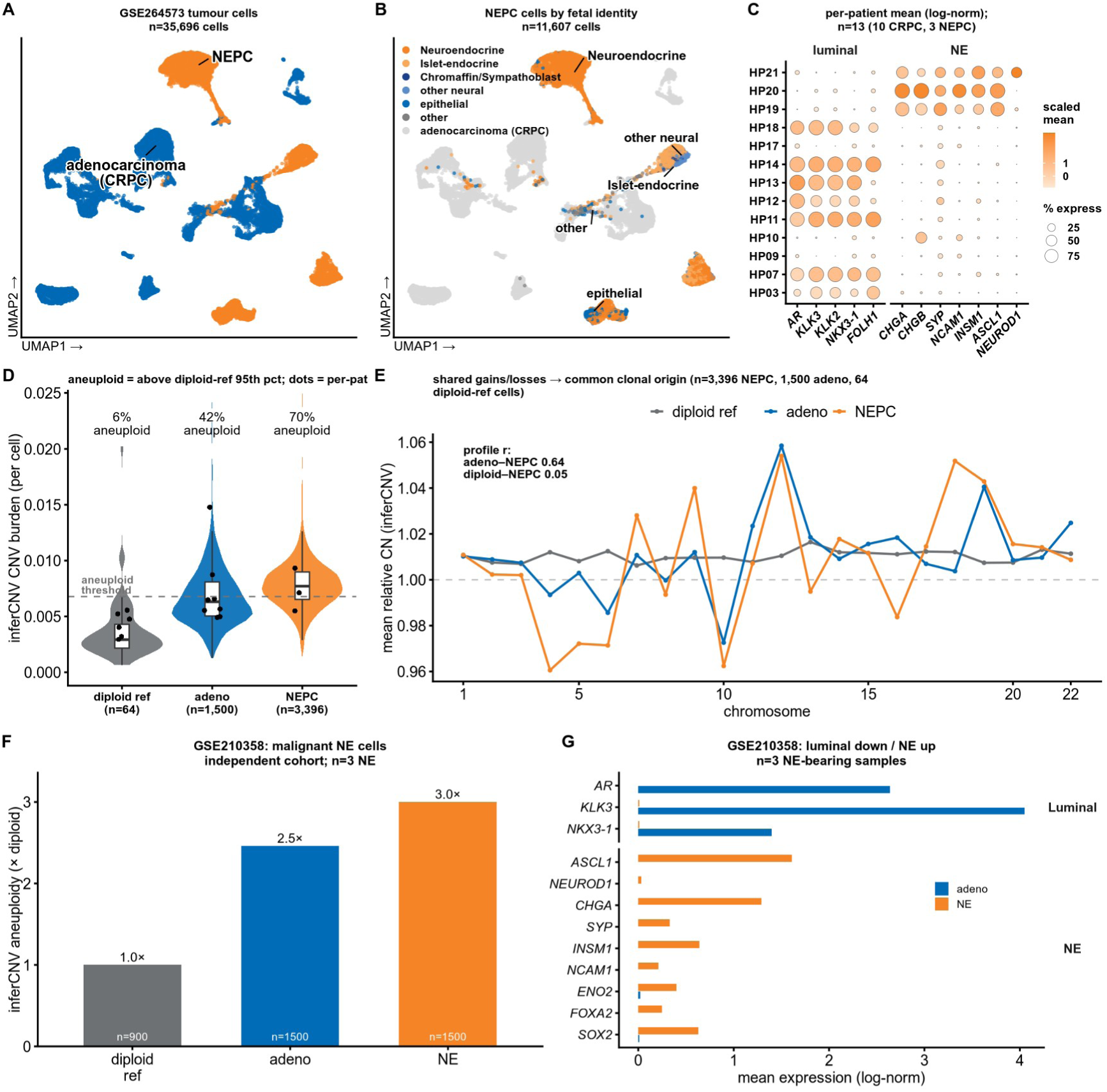
NEPC malignant cells share copy-number architecture with adenocarcinoma and show a luminal-to-neuroendocrine shift. **(A)** UMAP of 35,696 tumor cells (GSE264573; 20 patients [17 CRPC, 3 NEPC]; Seurat mutual-nearest-neighbor (MNN) integration) by transcriptional status; NEPC *n* = 11,607, adenocarcinoma/CRPC *n* = 24,089. Per-patient analyses use the 13 patients with ≥ 100 cells. **(B)** NEPC cells (*n* = 11,607) colored by argmax fetal cell type (Descartes atlas; rank correlation, no rejection). **(C)** Per-patient mean expression of luminal (*AR, KLK3, KLK2, NKX3-1, FOLH1*) and neuroendocrine (*CHGA, CHGB, SYP, NCAM1, INSM1, ASCL1, NEUROD1*) markers; 13 patients (10 CRPC, 3 NEPC). Fill = z-score across patients per gene; dot size = % expressing. **(D)** Per-cell inferCNV burden for the diploid reference (immune/endothelial, *n* = 64), adenocarcinoma (*n* = 1,500) and NEPC (*n* = 3,396). Violin = per-cell; box = median/interquartile range (IQR); dots = per-patient means; dashed line = aneuploidy threshold (95th percentile of the reference); % aneuploid per group. **(E)** Mean relative copy number per chromosome (1–22) for the three groups. Pearson *r*: adenocarcinoma–NEPC = 0.64; diploid-reference–NEPC = 0.05. **(F)** inferCNV aneuploidy (fold over diploid reference), GSE210358 (3 samples with NE features): reference 1.0× (*n* = 900), adenocarcinoma 2.5× (*n* = 1,500), NE 3.0× (*n* = 1,500). **(G)** Mean expression of luminal (*AR, KLK3, NKX3-1*) and neuroendocrine (*ASCL1, NEUROD1, CHGA, SYP, INSM1, NCAM1, ENO2, FOXA2, SOX2*) genes, NE vs adenocarcinoma (GSE210358).

To test whether NEPC cells were malignant, per-cell copy-number variation (CNV) was inferred with inferCNV against an internal diploid reference of immune and endothelial cells (n = 64 cells). NEPC and adenocarcinoma cells were more aneuploid than the reference (70% and 42% vs 6%; Fig. 1D). At the chromosome level, mean copy-number profiles correlated between adenocarcinoma and NEPC (Pearson *r* = 0.64) but not with the diploid reference (*r* = 0.05; Fig. 1E), so NEPC shared copy-number architecture with adenocarcinoma. Because inferCNV reports relative copy number without allele information, this is consistent with clonal relatedness but does not prove it.

The pattern reproduced in an independent single-cell cohort (three samples with neuroendocrine (NE) features, exploratory): NE cells were more aneuploid than adenocarcinoma and the diploid reference (3.0× vs 2.5× vs 1.0×; Fig. 1F), with luminal marker loss and neuroendocrine marker gain (Fig. 1G).

The signal was unlikely to be technical: scDblFinder doublet rates were <10% in every NEPC patient (5.0–9.0%; Fig. S1C), and the malignancy call used an internal diploid reference and a reference-derived aneuploidy threshold (Fig. 1D). These data support a malignant NEPC population that shares copy-number architecture with adenocarcinoma and shows a luminal-to-neuroendocrine shift, not an independent non-malignant neuroendocrine population; they do not yet resolve which neuroendocrine lineage it most resembles.

### NEPC cells exhibit a fetal neuroendocrine transcriptional program

We next asked which developmental neuroendocrine identity this state resembled. Transcription factor activity from CollecTRI regulons (decoupleR univariate linear model, ULM) separated NEPC from adenocarcinoma: ASCL1, NEUROD1 and MYCN were higher and REST and AR lower in NEPC (n = 3 NEPC vs 10 adenocarcinoma patients; Fig. 2A), with the same per-patient pattern across 13 patients for 12 lineage transcription factors (Fig. 2B). ASCL1, REST and AR exceeded size-matched random regulon nulls (permutation *P* = 0.005; NEUROD1 *P* = 0.025; Fig. 2A, Fig. S1E), so the signal was regulon-specific.

**Figure 2.**
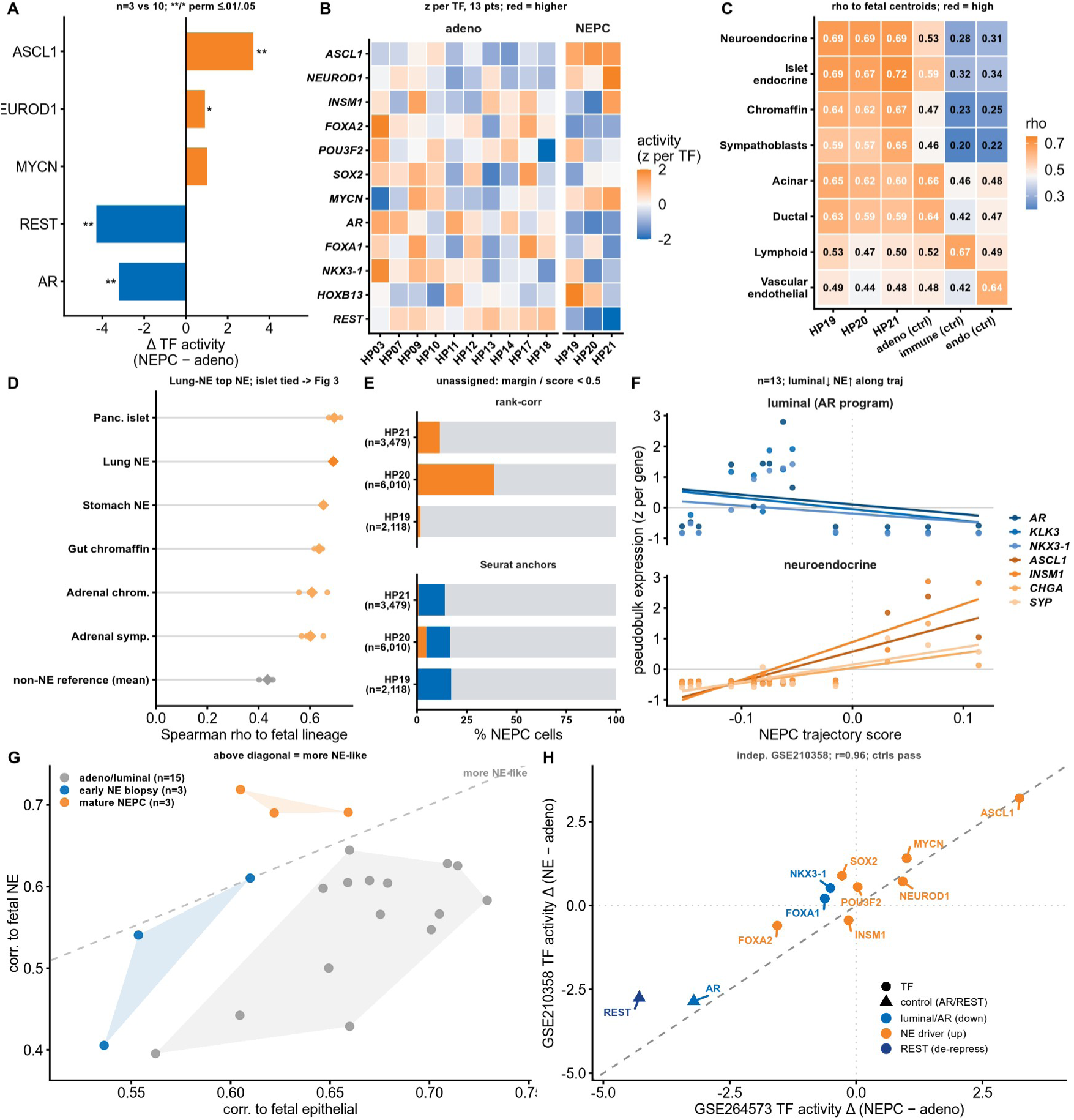
NEPC cells exhibit a fetal neuroendocrine transcriptional program. **(A)** Differential transcription factor (TF) activity for five lineage TFs (decoupleR univariate linear model [ULM], CollecTRI; Δ = mean NEPC − adenocarcinoma); *n* = 3 NEPC vs 10 adenocarcinoma patients. Double and single asterisks indicate permutation *P* ≤ 0.01 and *P* ≤ 0.05, respectively, versus size-matched random regulons (*ASCL1*, *REST*, *AR P* = 0.005; *NEUROD1 P* = 0.025); *MYCN* not tested. **(B)** Per-patient TF activity (decoupleR ULM, CollecTRI) for 12 TFs across 13 patients (10 adenocarcinoma, 3 NEPC). Fill = z-score across patients per TF. **(C)** Pseudobulk Spearman ρ of the three NEPC samples (HP19/20/21) and three controls (adenocarcinoma, immune, endothelial) to Descartes fetal cell-type centroids (172 organ–cell-type profiles collapsed to cell type by the max across organs). A cross-tissue mapping; the organ identity of the best match is resolved in (D) and Fig. 3. **(D)** Organ-resolved counterpart of (C): pseudobulk ρ to organ-specific fetal neuroendocrine/endocrine lineages plus a non-neuroendocrine reference (immune/endothelial/stromal mean). Dots = 3 NEPC patients; diamond = lineage mean. Lung-neuroendocrine and pancreatic islet-endocrine are tied (both ρ = 0.69; islet slightly higher before rounding), motivating the lung versus islet checks in Fig. 3. **(E)** Per-cell reference mapping of NEPC cells (HP19 *n* = 2,118; HP20 *n* = 6,010; HP21 *n* = 3,479) by two methods (rank correlation; Seurat anchor transfer). Bars = % cells confidently assigned a neuroendocrine-family label (orange), another label (blue), or unassigned (grey; margin or prediction score < 0.5). **(F)** Per-patient pseudobulk expression (z-score per gene) of luminal (*AR, KLK3, NKX3-1*) and neuroendocrine (*ASCL1, INSM1, CHGA, SYP*) genes vs the NEPC trajectory score; *n* = 13 patients; lines = per-gene linear fits. **(G)** Per-sample pseudobulk ρ to fetal epithelial (*x*) and fetal neuroendocrine (*y*) centroids for adenocarcinoma/luminal (*n* = 15), early-NE (*n* = 3) and mature-NEPC (*n* = 3) samples. Dashed line = identity; shaded = per-group convex hulls. **(H)** Differential TF activity (decoupleR ULM, CollecTRI, pseudobulk), GSE264573 (Δ NEPC − adenocarcinoma, *x*) vs GSE210358 (Δ NE − adenocarcinoma, *y*) for 11 lineage TFs; triangles = *AR*/*REST* controls. Pearson *r* = 0.96, Spearman ρ = 0.89; controls pass; 7/11 TFs agree in direction. GSE210358 is sample-level, exploratory.

Pseudobulk profiles of the three NEPC samples were then correlated (Spearman) with Descartes fetal cell-type centroids. NEPC was most similar to neuroendocrine and endocrine fetal types; the adenocarcinoma, immune and endothelial controls were not (Fig. 2C). Resolved by organ, fetal lung-neuroendocrine and pancreatic islet-endocrine reached nearly identical scores (both ρ = 0.69; Fig. 2D), so the organ-level assignment was unresolved at this stage.

Per-cell mapping by two independent methods (rank correlation and Seurat anchor transfer) gave NEPC cells a neuroendocrine-family fetal label in all three patients, leaving low-confidence cells unassigned (Fig. 2E). The reference spanned all 77 fetal cell types (open-world; Fig. S1B), and MetaNeighbor, which misplaced the controls, was excluded (Fig. S1A). The high unassigned fraction and open-world reference argued against a closed-world artifact.

Along a per-patient NEPC trajectory score, luminal genes fell and neuroendocrine genes rose (n = 13 patients; Fig. 2F), and samples ordered from adenocarcinoma through early-NE to mature-NEPC on epithelial-versus-neuroendocrine axes (n = 15 / 3 / 3 samples; Fig. 2G). The program reproduced in the same independent single-cell cohort (Pearson *r* = 0.96, Spearman ρ = 0.89; exploratory, three-sample), with AR and REST controls in the expected direction (Fig. 2H).

NEPC cells thus showed a fetal neuroendocrine program, though the data did not separate a pulmonary from an islet-endocrine identity.

### Subtype-resolved mapping supports a lineage-restricted fetal pulmonary neuroendocrine-like identity

Because NEPC arises by dedifferentiation and lineage plasticity, we compared it against fetal rather than adult developmental references, consistent with oncofetal reprogramming as a route to plasticity [11,12]. To distinguish a specific fetal neuroendocrine identity from a generic one, NEPC cells were scored on a fetal-minus-adult lung-neuroendocrine program (controlled using T cells for the generic fetal-to-adult shift). The program was higher in NEPC than adenocarcinoma (two-sided Mann–Whitney *P* = 0.008, Cliff’s δ = +1.0; n = 3 NEPC vs 7 adenocarcinoma patients; Fig. 3A), and the effect was consistent in direction across three independent cohorts (per-cohort Hedges’ *g* = 2.45, 1.64 and 1.45; random-effects pooled *g* = 1.62, 95% CI 1.18–2.06; Fig. 3B); with only three cohorts *I*² and the pooled interval are imprecise, so we read this as directional replication rather than a precise pooled estimate.

**Figure 3.**
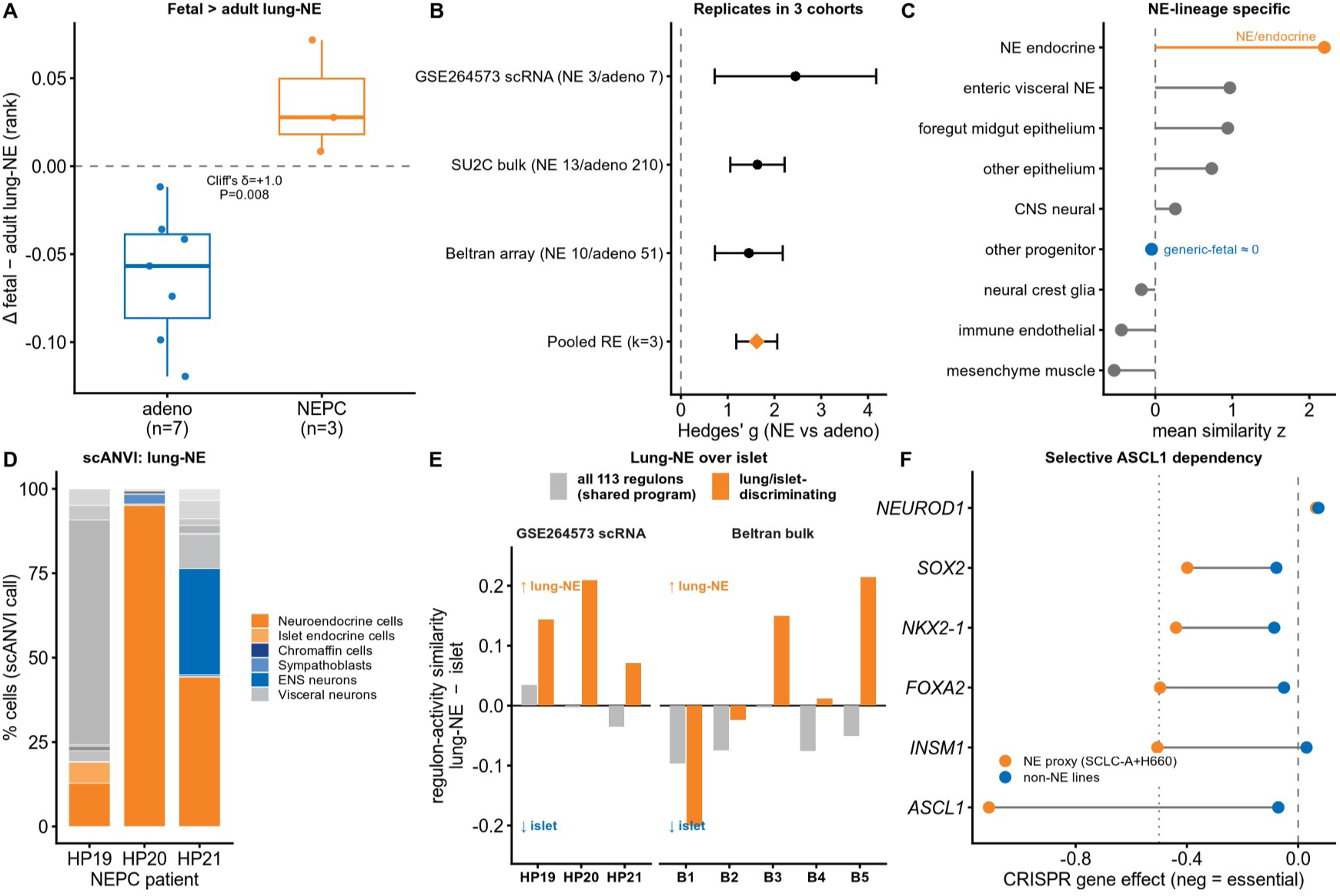
Subtype-resolved mapping supports a lineage-restricted fetal pulmonary neuroendocrine-like identity. **(A)** Fetal-minus-adult lung-NE program score (rank percentile) per GSE264573 sample; each fetal/adult program scored independently per sample, with the generic fetal-vs-adult/platform shift removed using a T cell control. Box = median ± IQR; points = patients. NEPC (*n* = 3, median Δ = +0.028) vs adenocarcinoma (*n* = 7, median Δ = −0.057); two-sided Mann–Whitney *P* = 0.008, Cliff’s δ = +1.0. (Adeno *n* = 7: this scoring’s ≥ 50-cell query keeps fewer patients than the 10 used elsewhere.) **(B)** Random-effects meta-analysis with metafor and restricted maximum likelihood (REML) of the NE-vs-adenocarcinoma effect on this axis across three cohorts: GSE264573 single-cell RNA-seq (scRNA-seq; 3 / 7), SU2C/WCDT bulk small-cell vs adenocarcinoma (13 / 210), Beltran array SYP⁺ vs CHGA⁻ (10 / 51). Per-cohort Hedges’ *g* = 2.45 / 1.64 / 1.45; pooled *g* = 1.62 (95% CI 1.18–2.06), *I²* = 0% (uninterpretable at k = 3 cohorts), *P* = 4.1 × 10⁻¹³. **(C)** Lineage specificity: mean similarity z-scored within each query (Spearman to the Descartes atlas with 77 cell types) of NEPC cells, collapsed into nine fetal families. NE/endocrine is highest (mean z approximately +2.2); generic progenitors are approximately 0. One-sided Wilcoxon over the 77 types, per patient, *P* = 1.1 × 10⁻⁸ / 1.4 × 10⁻⁹ / 4.9 × 10⁻⁶ (HP19/20/21), all < 1.5 × 10⁻⁵ after Bonferroni. Foregut-midgut epithelium is also elevated. **(D)** Batch-aware mapper: per-patient scANVI label fractions onto the fetal Descartes reference; maximum posterior < 0.5 = uncertain/decoy. Confident NE-family fraction HP19/20/21 = 22 / 98 / 86%. Within all scANVI calls the fetal Lung-NE label accounted for 12.8 / 95.0 / 44.2% (largest confident NE-family component); neural crest Chromaffin was approximately 0% (0.2 / 0.1 / 0.0%). HP19 is predominantly decoy. **(E)** Regulon-activity test of the islet competitor: NEPC similarity to fetal lung-NE minus islet, on all 113 shared regulons (grey) and the 16 regulons selected from the reference to discriminate lung NE from islet (orange; query-blind), per scRNA patient (HP19–21) and Beltran sample (B1–B5). On the discriminating regulons NEPC favors lung-NE in 3/3 scRNA and 3/5 bulk (robust to dropping FOXA1); on all 113 the two are tied. Caveat: the lung NE versus islet distinction is sensitive to method choice and associated with FOXA2; this is a lean, not a resolved separation. **(F)** Selective lineage-TF dependency (DepMap 24Q4, Chronos) in the malignant NE-lineage proxy (ASCL1-high SCLC-A lines + NEPC line NCI-H660) vs non-NE lines; no fetal PNEC line exists in DepMap. ASCL1 is selectively essential (effect −1.11 vs −0.07; Δ = −1.04; selective dependency rank 2 / 17,916; *P* = 8.3 × 10⁻⁹); INSM1/FOXA2/NKX2-1/SOX2 also selective (rank 11 / 19 / 39 / 57); NEUROD1 is not (rank 7,585; within-panel specificity control). Dependency shown in the proxy, not NEPC tumors. *Effective n = patients/samples, not cells: 3 NEPC scRNA patients (HP19–21) underlie A, C, D and the scRNA arm of E; bulk cohorts add samples in B and E*.

The resemblance was lineage-restricted: within the query, similarity to the fetal atlas with 77 cell types was highest for the neuroendocrine/endocrine family (mean z approximately 2.2) and near zero for generic progenitors, ranking highest in every patient (one-sided Wilcoxon over the 77 types, all *P* < 1.5 × 10⁻⁵ after Bonferroni; Fig. 3C). A batch-aware mapper (scANVI) assigned 22%, 98% and 86% of cells to the neuroendocrine family across the three patients (seed-42 reference run; Fig. 3D), within which fetal lung-neuroendocrine was the largest subtype in HP20 and HP21 and chromaffin was near zero. Across five random seeds these fractions were robust for HP20 (median lung-neuroendocrine 95%) and HP21 (35%) but unstable for HP19 (lung-neuroendocrine 1–51%; confident neuroendocrine 10–68%), consistent with its decoy-heavy profile (per-seed distributions, Fig. S2G; an earlier pipeline run, Fig. S2D–E). The per-patient identity assignment therefore rests on the deterministic rank-correlation and anchor-transfer mappings (Fig. 3C, 3E), which do not depend on initialization.

Rank-based subtype mapping placed fetal lung-neuroendocrine as the top match in all eight NEPC samples (Fig. S2A), and alternative lineages were excluded directly: lung specifiers NKX2-1/TTF1 were detected in 16–78% of NEPC cells versus 4% of adenocarcinoma, whereas islet (PDX1/NKX6-1) and sympathoblast (PHOX2B/HAND2/GATA3) specifiers were near zero (Fig. S2B). The mapper recovered known external references and re-recovered held-out fetal cells (Fig. S2C,F): neuroblastoma — a neuroendocrine, ASCL1-expressing tumor of neural-crest origin — mapped to chromaffin and other neural-crest lineages (0/36 to pulmonary neuroendocrine), whereas SCLC-A (29/30) and NEPC (3/3) mapped to pulmonary neuroendocrine, so the pulmonary-neuroendocrine assignment was lineage-specific, not a default for any neuroendocrine or ASCL1-driven state. On the 16 reference-defined regulons that best separate lung-NE from islet, NEPC leaned toward lung-NE in 3/3 single-cell and 3/5 bulk samples, though the two were tied on all 113 shared regulons (Fig. 3E); a de novo regulon-activity space likewise placed NEPC with the endocrine (lung-NE plus islet) pair under discriminability and permutation controls (Fig. S3).

In DepMap, ASCL1 was the strongest selective dependency among the neuroendocrine regulators tested (gene effect −1.11 vs −0.07; rank 2 of 17,916; *P* = 8.3 × 10⁻⁹), with INSM1, FOXA2, NKX2-1 and SOX2 also selective and NEUROD1 not (Fig. 3F); this used a malignant neuroendocrine proxy, not NEPC tumors. Taken together, a fetal pulmonary neuroendocrine-like assignment was favored over islet, chromaffin and sympathoblast alternatives, with the distinction between lung and islet remaining sensitive to method choice and associated with FOXA2.

### The luminal-to-neuroendocrine switch follows an ordered transcriptional sequence that is consistent across cohorts

The luminal-to-neuroendocrine transition resolved into an ordered gene set grouped by role (luminal, repressor, neuroendocrine driver, target and effector nodes), scored by rank per patient (Fig. 4A). Per-patient pseudobulk differential expression (DESeq2; n = 3 NEPC vs 10 adenocarcinoma patients) showed loss of luminal markers and gain of neuroendocrine drivers and effectors (Fig. 4B): NKX3-1 log₂FC = −9.39, ASCL1 = +8.42, CHGA = +7.22, INSM1 = +5.50, SYP = +2.66 and REST = −2.46 (all false discovery rate [FDR] < 0.05; NEUROD1 below the count threshold). Because this is a shift across the lineage, FDR < 0.05 spanned 11% of the genome, so the shrunken effect size was more informative than the adjusted *P* value.

**Figure 4.**
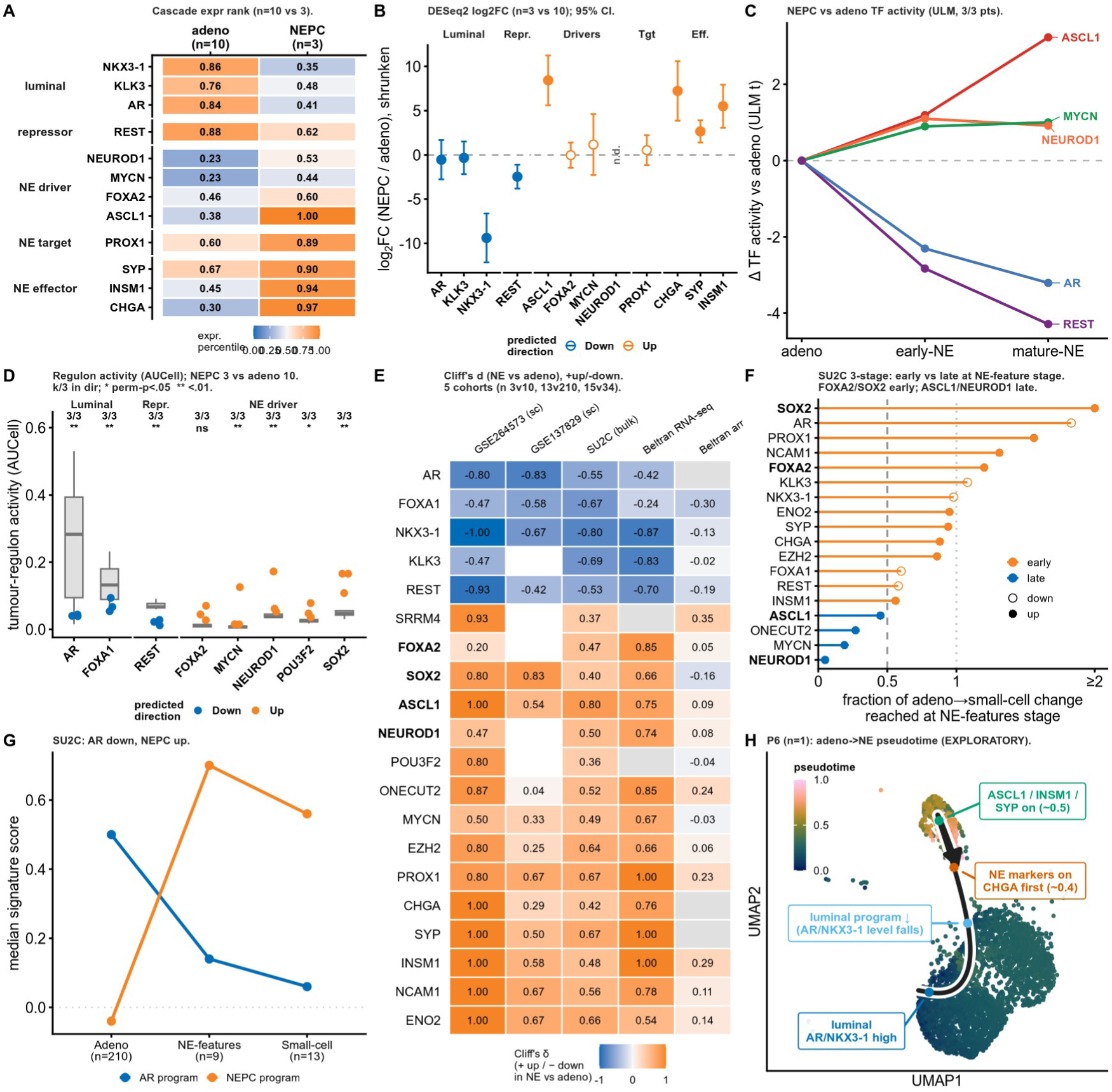
The luminal-to-neuroendocrine switch follows an ordered transcriptional sequence consistent across cohorts. **(A)** Expression-percentile heatmap of the 12-gene luminal-to-NE cascade in GSE264573, adenocarcinoma (n = 10) vs NEPC (n = 3, HP19–21), per patient, scored by rank; rows grouped by role. Luminal markers (AR, KLK3, NKX3-1) and the repressor REST fall; NE drivers (ASCL1, FOXA2, NEUROD1, MYCN), the target PROX1 and effectors (CHGA, SYP, INSM1) rise. **(B)** Per-patient pseudobulk DESeq2 (GSE264573, raw counts), NEPC (n = 3) vs adenocarcinoma (n = 10); apeglm-shrunken log₂FC ± 95% CI; filled = FDR < 0.05, n.d. = untestable (NEUROD1). Direction agrees for 10/12 switches; strong nodes: NKX3-1 −9.39, ASCL1 +8.42, CHGA +7.22, INSM1 +5.50, SYP +2.66, REST −2.46 (all FDR < 0.05). The shift across the lineage spans 11% of the genome at FDR < 0.05, so the shrunken effect size, not padj, is informative. **(C)** decoupleR ULM TF activity (CollecTRI), Δ vs each dataset’s own adenocarcinoma, across the adeno, early-NE and mature-NE stages (GSE264573 = adeno + mature-NE endpoints; GSE137829 = early-NE). ASCL1, NEUROD1, MYCN rise and REST is de-repressed while AR falls; 3/3 mature-NE patients consistent. **(D)** De novo pySCENIC regulons (GRNBoost2 + cisTarget, built on GSE264573), AUCell-scored per patient, NEPC (n = 3) vs adenocarcinoma (n = 10); k/3 = patients in the predicted direction; double and single asterisks indicate the shuffled-gene permutation null at P < 0.01 and P < 0.05, respectively (100 permutations, minimum P approximately 0.0099). All eight programs were consistent in 3/3 patients: luminal AR/FOXA1 and REST fall, whereas NE drivers SOX2/NEUROD1/MYCN rise; POU3F2 is weaker and FOXA2 is not significant. AR-down is the positive control. **(E)** Cliff’s δ (NE vs adenocarcinoma) for 20 lineage genes across five datasets (four cohorts; Beltran contributes both RNA-seq and array): GSE264573 scRNA (3 / 10), GSE137829 scRNA (6 / 4 per-patient pseudobulks from 6 patients), SU2C/WCDT bulk (13 / 210), Beltran RNA-seq (15 / 34), Beltran array (20 / 51). Direction is consistent and matches the NE prediction for nearly every gene except in the array: 19/20 genes (GSE264573 scRNA) and all 18 testable (Beltran RNA-seq) vs 13/17 (array, which floors low abundance NE-TFs). SRRM4 is the lone recurrent disagreer. **(F)** Switch ordering: SU2C/WCDT three-stage series, adenocarcinoma (n = 210), adeno with NE features (n = 9) and small-cell (n = 13). Per gene, the fraction of the total adeno-to-small-cell change reached at the intermediate stage (≥ 0.5 = early, < 0.5 = late); Jonckheere increasing-trend *P* ≤ 0.002 (Benjamini-Hochberg) for the 18 orderable genes. FOXA2/SOX2 (and AR loss) early; ASCL1/NEUROD1/MYCN late. (20 evaluated; SRRM4 and POU3F2 untimable, not shown.) A relative ordering across morphological stages, not real time. **(G)** Stage-axis validity: SU2C/WCDT AR-program and NEPC-program scores (median per stage; n = 210 / 9 / 13 samples). AR program 0.50, 0.14, 0.06; NEPC program −0.04, 0.70, 0.56 across the three stages, supporting the stage axis as a luminal-to-NE axis. **(H)** Single-tumor Slingshot pseudotime UMAP (GSE137829 patient P6; n = 1 tumor): luminal AR/NKX3-1 high, then luminal-program loss, then CHGA onset, then ASCL1/INSM1/SYP onset (FOXA2/NEUROD1 unmeasured here). Exploratory; pseudotime is not real time and illustrates, rather than establishes, the A–G ordering. *Effective n = patients/samples, not cells: GSE264573 per-patient pseudobulk underlies A–D; bulk/array and the three-stage series add samples in E–G; H is one tumor*.

Transcription factor and regulon activity tracked the same direction. decoupleR activity rose for ASCL1, NEUROD1 and MYCN and fell for AR and REST across the adenocarcinoma, early-NE and mature-NE stages (Fig. 4C; per-stage detail in Fig. S4A); the early-NE stage came from an incompletely transdifferentiated cohort whose provided neuroendocrine label over-calls committed cells approximately 3-fold (Fig. S1D). De novo pySCENIC regulons moved consistently in all three NEPC patients across eight programs: the neuroendocrine drivers SOX2, NEUROD1 and MYCN rose and the luminal and REST regulons fell (permutation *P* < 0.01), while POU3F2 was weaker and FOXA2 not significant (Fig. 4D).

The direction reproduced in every cohort. Cliff’s δ matched the neuroendocrine prediction for 19 of 20 genes in the discovery single-cell cohort and all 18 testable genes in Beltran RNA-seq, with weaker agreement in the low abundance microarray (13/17), across five datasets (Fig. 4E); SRRM4 was the only recurrent disagreer. In the SU2C/WCDT three-stage morphological series (n = 210 / 9 / 13 samples), the AR program fell from 0.50 to 0.14 to 0.06 and the NEPC program rose from −0.04 to 0.70 to 0.56 (Fig. 4G). The 18 orderable genes showed a Jonckheere trend (*P* ≤ 0.002), FOXA2 and SOX2 changing early and ASCL1, NEUROD1 and MYCN late (Fig. 4F). A composite that treated samples as stages rose monotonically (Jonckheere *P* < 0.001, n = 21 samples; Fig. S4B,D), and the strong cascade nodes passed the adenocarcinoma range in all three NEPC patients (Fig. S4C).

A single transdifferentiating tumor (patient P6 of that transdifferentiation cohort) showed the same ordering along Slingshot pseudotime, from luminal-high to neuroendocrine marker onset (Fig. 4H); this panel is exploratory (n = 1 tumor from 1 patient) and pseudotime is not elapsed time. The sequence was therefore ordered and reproducible in all cohorts, not a timed developmental program.

### An unbiased regulatory screen nominates a neuroendocrine circuit centered on ASCL1 with FOXA1 binding

To find candidate regulators without pre-selection, we examined genome-wide differential expression first (n = 3 NEPC vs 10 adenocarcinoma patients; 24,828 genes; 1,111 up and 1,508 down at FDR < 0.05). Neuroendocrine markers rose (ASCL1 log₂FC = +8.4, CHGA +7.2, INSM1 +5.5, DLL3 +5.4, SYP +2.7) and the luminal/AR program fell (NKX3-1 −9.4, REST −2.5, AR −0.5; Fig. 5A). Several nominated regulators were not recoverable by fold change alone (SOX2 +0.38; POU3F2 +0.07, not significant; NEUROD1 below bulk detection), motivating inference across methods.

**Figure 5.**
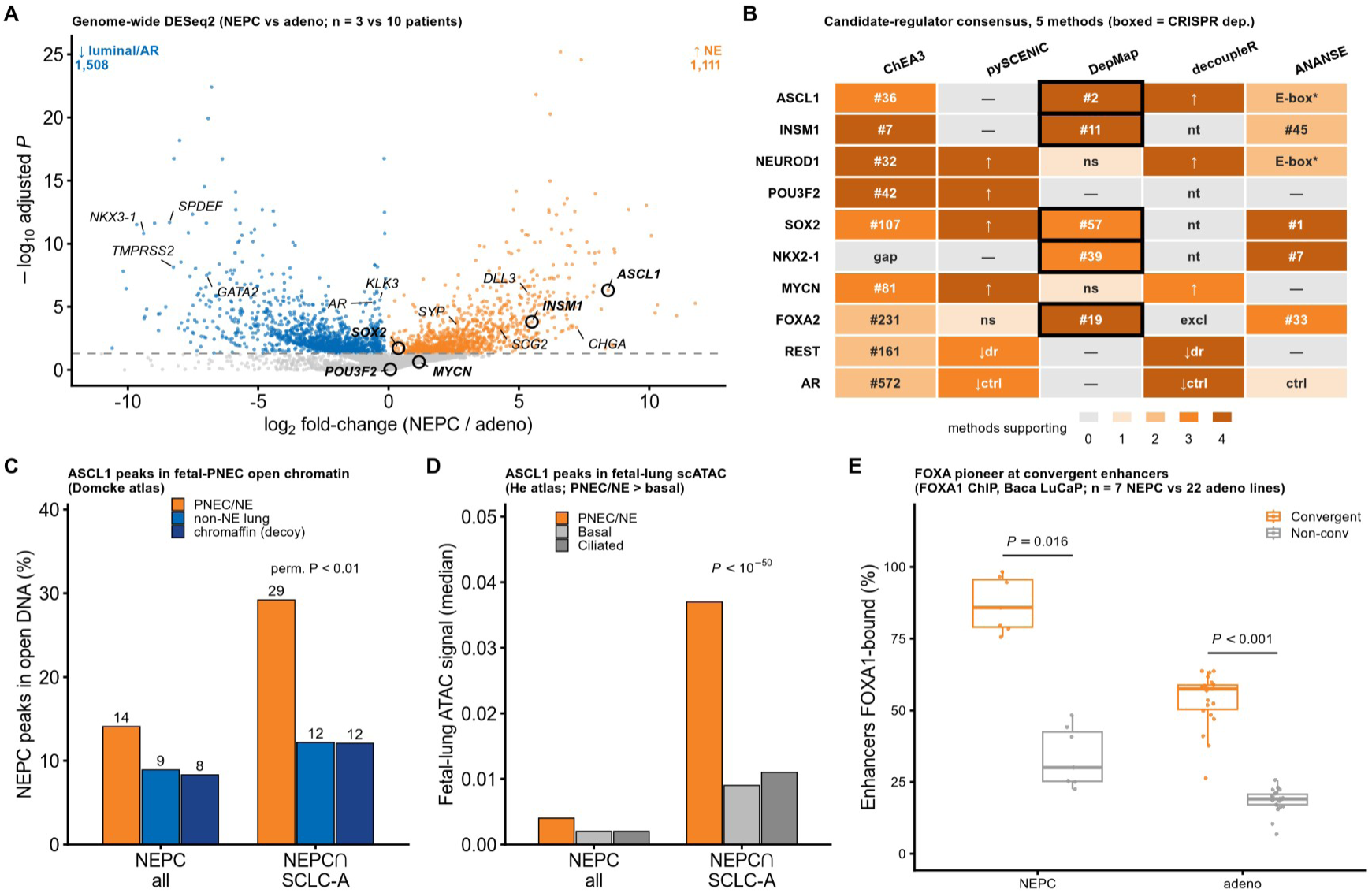
An unbiased regulatory screen nominates a neuroendocrine circuit centered on ASCL1 with FOXA1 binding. **(A)** Volcano of genome-wide pseudobulk DE (per-patient aggregated counts), NEPC (n = 3 patients, HP19–21) vs adenocarcinoma (n = 10 patients), GSE264573; 24,828 genes, 1,111 up / 1,508 down at FDR < 0.05. NE markers rise (ASCL1 +8.4, CHGA +7.2, INSM1 +5.5, DLL3 +5.4, SCG2 +4.2, SYP +2.7) and the luminal/AR program falls (NKX3-1 −9.4, SPDEF −8.4, TMPRSS2 −8.3, GATA2 −7.0, REST −2.5, AR −0.5). Several nominated TFs are not findable by fold change (SOX2 +0.38; POU3F2 +0.07, NS; NEUROD1 below detection), motivating (B). **(B)** Candidate regulator consensus across five orthogonal methods. Rows = candidate transcription factors; fill = number of the four inference methods (ChEA3 target enrichment, pySCENIC and decoupleR regulon activity, ANANSE enhancer network influence) supporting each (0–4); black-outlined cells mark a selective DepMap CRISPR dependency. The convergent read-out is a neuroendocrine transcription factor circuit centered on ASCL1 with INSM1, NEUROD1, POU3F2 and SOX2; ASCL1 is the apex node (DepMap selective dependency rank 2/17,916). FOXA1 is a constitutive pioneer (baseMean approximately 5,000; log₂FC = −0.7, not significant), so it is absent from this differential-expression consensus; its role is occupancy/redeployment (panel E), and FOXA1 and FOXA2 share the FOXA motif (Fig. S5D). Full per-method rankings are provided in Source Data. **(C)** Convergent ASCL1 cistrome vs Domcke fetal open chromatin: fraction of NEPC ASCL1 peaks (5,531) and the shared NEPC/SCLC-A subset (1,210) in pulmonary neuroendocrine (14% / 29%) over non-NE lung (9% / 12%) or a chromaffin decoy (8% / 12%); permutation z = 102 / 95. **(D)** Replication in a second atlas: median fetal lung scATAC signal (He et al. [18]) at the same peaks; PNEC/NE exceeds basal (P < 10⁻⁵⁰ for both shared NEPC/SCLC-A and NEPC-all). Cell line cistromes; effective patient n in (A). **(E)** FOXA1 binds the convergent enhancers (Baca GSE161948 LuCaP ChIP, hg19): FOXA1 occupies the convergent set (CONV, 353 enhancers) far more than non-convergent ASCL1 peaks: 87.0% vs 33.8% in NEPC (7 lines; paired Wilcoxon P = 0.016) and 53.8% vs 18.5% in adenocarcinoma (22 lines; P < 0.001). Chromatin-level validation (C–E) covers ASCL1 and FOXA1 only; the other nominated TFs rest on the method consensus and CRISPR dependency evidence in (B). Cistrome and ChIP are cell line/PDX, not patient NEPC.

Five orthogonal methods (ChEA3 target enrichment, pySCENIC and decoupleR regulon activity, DepMap dependency, ANANSE enhancer network influence) converged on a neuroendocrine transcription factor circuit centered on ASCL1 with INSM1, NEUROD1, POU3F2 and SOX2 (Fig. 5B); the ANANSE model also recovered the AR/luminal program as a positive control and nominated SOX2 and NKX2-1 for the transition (Fig. S5F). FOXA1 itself was not differentially expressed (baseMean approximately 5,000; log₂FC = −0.7, not significant) and so did not surface in this screen.

The convergent regulatory elements were neuroendocrine lineage chromatin. The ASCL1 cistrome (5,531 peaks) and its shared NEPC/SCLC-A subset (1,210 peaks) fell preferentially in fetal pulmonary neuroendocrine open chromatin of the Domcke atlas (14% and 29%; Fig. 5C), and less in non-neuroendocrine lung epithelium (9% and 12%) or a chromaffin decoy (8% and 12%; permutation z = 102 / 95). This reproduced in an independent fetal lung single-cell assay for transposase-accessible chromatin (scATAC) atlas (pulmonary neuroendocrine above basal, *P* < 10⁻⁵⁰; Fig. 5D). Lineage and enhancer specificity, and the single passing NEPC ASCL1 cistrome used for the motif gate, are in Fig. S5A–C.

FOXA1 occupied these elements directly. In deposited chromatin immunoprecipitation (ChIP), FOXA1 bound the convergent enhancers (353 elements) more often than non-convergent ASCL1 peaks, in NEPC (87.0% vs 33.8%; 7 cell lines; *P* = 0.016) and in adenocarcinoma (53.8% vs 18.5%; 22 cell lines; *P* < 0.001; Fig. 5E), consistent with the FOXA motif (Fig. S5D) and H3K27ac activity (Fig. S5E) at the same elements (paired test across LuCaP lines; Methods). These analyses nominate, but do not prove, an architecture centered on ASCL1 with FOXA1 binding: the evidence is associative (n = 3 NEPC patients), with the ASCL1 cistrome from cell line/SCLC-A models and the FOXA1/H3K27ac ChIP from LuCaP lines rather than primary patient NEPC.

### Spatial transcriptomics localizes the fetal pulmonary neuroendocrine-like program to malignant NEPC territory

We then examined the program in tissue. In a NEPC Visium section (one section from one patient), a fetal pulmonary neuroendocrine marker program colocalized with a pan-neuroendocrine program (per-spot Spearman ρ = +0.55) but not with a Club cell control (−0.51; Fig. 6A–B), and it discriminated the malignant region defined by copy number (AUROC 0.83) above an AR/luminal set (0.56) and the Club control (0.20). The discrimination held after residualizing for spot complexity (AUROC 0.85; permutation *P* = 0.005; 531 malignant vs 1,119 non-malignant spots; Fig. 6C).

**Figure 6.**
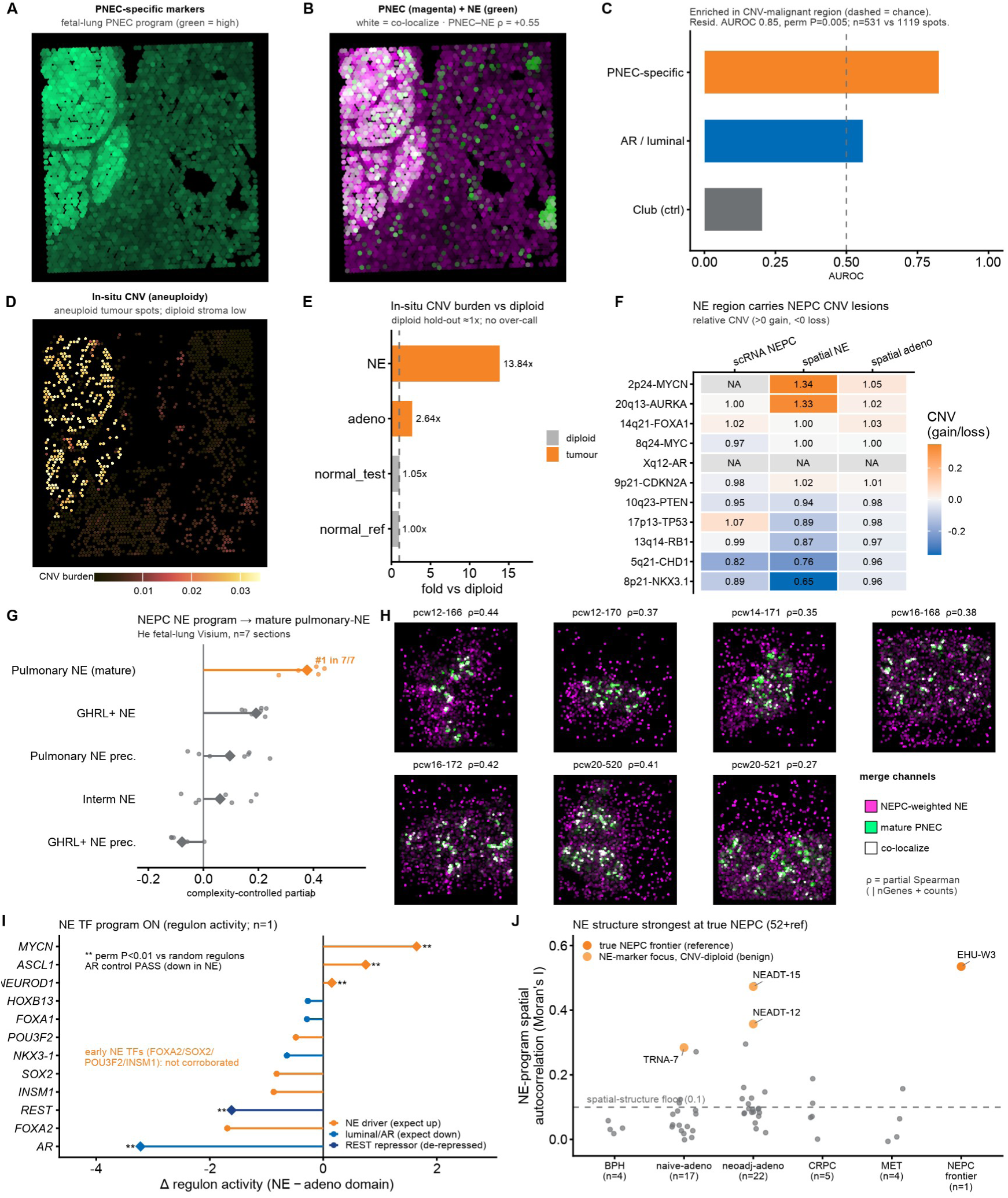
Spatial transcriptomics localizes the fetal pulmonary neuroendocrine-like program to malignant NEPC territory. **(A–C) Forward analysis: the fetal lung PNEC program concentrates in the malignant NE region of human NEPC tissue Visium**, GSE230282 EHU-W3; one section from one patient). **(A)** Spatial map of PNEC-specific marker-program activity (green = high). **(B)** Two-channel overlay of the PNEC (magenta) and pan-NE (green) programs; white = colocalization. Per-spot Spearman ρ = +0.55 (PNEC vs NE) versus −0.51 for the Club cell negative control. **(C)** Specificity: AUROC for each gene set being higher inside the malignant region defined by CNV than outside. The fetal lung PNEC set (orange) discriminates the malignant region (raw AUROC = 0.83), far above the AR/luminal set (0.56) and the Club control (0.20); dashed line = chance. The permutation P belongs to the complexity-residualized analysis: residualized AUROC = 0.85, permutation P = 0.005 (531 malignant vs 1119 non-malignant spots). **(D–F) In situ analysis: the NE region is genuine aneuploid tumor carrying canonical NEPC copy-number lesions** (inferCNV, same section / one patient). **(D)** Spatial CNV-burden map (aneuploid tumor spots high; diploid stroma low). **(E)** CNV burden relative to a diploid baseline: NE spots 13.8×, adeno spots 2.6×; a held-out diploid test returns 1.05× (control; the method is not over-calling). **(F)** Driver-locus heatmap (relative copy number; > 0 gain, < 0 loss) across scRNA NEPC (GSE264573), spatial NE and spatial adeno. The in situ NE carries the textbook NEPC lesions in the same direction as independent scRNA NEPC (MYCN/AURKA gain; RB1/TP53/PTEN/NKX3-1/CHD1 loss; 8/10 concordant). **(G, H) Reverse analysis: the NEPC NE program localizes to the mature pulmonary-NE compartment of fetal lung** (He fetal lung Visium; 7 sections). **(G)** Five fetal lung NE substates ranked by the complexity-controlled partial Spearman correlation (controlling per-spot nGenes + counts) between the NE score weighted by the NEPC program and each substate’s cell2location abundance; one dot per section, diamond = mean. Mature “Pulmonary neuroendocrine” ranked first in 7/7 sections (mean partial ρ = 0.38; GHRL⁺ NE second at 0.19). **(H)** The same NE score weighted by the NEPC program (magenta) and mature-PNEC cell2location abundance (green) as a fluorescence-style overlay for all 7 sections (white = colocalization); per-section partial ρ annotated. The ranking is driven by the canonical NE gene set rather than weighting unique to NEPC (weight permutation P approximately 0.5) and is at the region level; it supports localization to the mature NE compartment of fetal lung, not specificity across organs (the latter is addressed by reference mapping in Figs 3–4). **(I, J) SPATIAL REGULATORY READOUTS. (I)** NE master TF regulon activity (decoupleR ULM on the CollecTRI network), NE minus adeno spatial domain (one section / one patient): NE drivers (MYCN, ASCL1, NEUROD1) up; the luminal/AR program and the REST repressor down (double asterisks indicate permutation P < 0.01 versus size-matched random regulons; the AR direction control passes). **(J)** Spatial autocorrelation (Moran’s I) of the NE program across 52 GSE278936 sections plus the EHU-W3 reference frontier. The EHU-W3 true NEPC frontier is the highest reference point; most staged sections away from the frontier are low, but several NEADT/TRNA sections show NE marker spatial structure and are labeled as CNV-diploid/benign foci in the source analysis. Dashed line = spatial structure floor 0.1. Spot/cell size per platform; n stated per panel (effective n = sections / patients). inferCNV reports relative copy number (no allele information).

That region was aneuploid tumor. Inferred copy-number burden was higher in neuroendocrine than adenocarcinoma spots or a held-out diploid test (13.8× vs 2.6× vs 1.05×; Fig. 6D–E), and the in situ neuroendocrine spots carried canonical NEPC lesions in the same direction as independent single-cell NEPC (8 of 10 driver loci concordant; Fig. 6F).

The reverse mapping placed the NEPC program in fetal lung. In seven fetal lung Visium sections, the neuroendocrine score weighted by the NEPC program correlated most strongly with the mature “pulmonary neuroendocrine” cell state, ranking first in 7 of 7 sections (mean complexity-controlled partial ρ = 0.38; GHRL⁺ neuroendocrine second at 0.19; Fig. 6G–H). This reflected the canonical neuroendocrine gene set rather than weighting unique to NEPC (weight permutation *P* approximately 0.5), and was at the region level, supporting localization within the lung to the mature compartment, not specificity between organs.

Neuroendocrine master-regulator activity was higher in the neuroendocrine than the adenocarcinoma spatial domain (MYCN, ASCL1 and NEUROD1 up; AR/luminal and REST down; permutation *P* < 0.01; Fig. 6I), and the program’s spatial structure was strongest at the true NEPC frontier across 52 sections of a NEPC foci Visium screen plus the EHU-W3 reference, with some neoadjuvant sections also structured (Fig. 6J). The in situ adenocarcinoma-to-neuroendocrine axis was spatially continuous for resolvable markers (KLK3 and CHGA Moran’s *I* = 0.759 and 0.619; Fig. S6A,C,D), whereas lineage transcription factors fell below the Visium detection floor and were spatially unorderable (Fig. S6B,C). In the negative control screen the real frontier had the highest CHGA spatial structure, though NEADT/TRNA sections were not all zero (Fig. S6E). These data place the fetal pulmonary neuroendocrine-like program within malignant NEPC territory while remaining exploratory at the section level (forward: n = 1 section from 1 patient; reverse: at the region level, not unique to NEPC).

## Discussion

NEPC emerges in advanced prostate adenocarcinoma under AR pathway inhibition and is clinically lethal [1,2]. That it carries a neuroendocrine program is established; what has been unclear is which neuroendocrine identity it represents. Here we refine the NEPC neuroendocrine state from a generic neuroendocrine phenotype to a fetal, lineage-restricted, pulmonary neuroendocrine-like state, aligned with an ordered transcriptional transition and associated with a regulatory architecture centered on ASCL1 with FOXA1 binding.

This conclusion rests on convergent evidence from single-cell, bulk, cistrome, and spatial data spanning multiple cohorts. The NEPC cells profiled here were malignant and shared copy-number architecture with the adenocarcinoma compartment in the discovery cohort, consistent with transdifferentiation from adenocarcinoma rather than an independent neuroendocrine population. Their transcriptional state was most similar to a fetal neuroendocrine program, was best matched among resolved lineages by a pulmonary neuroendocrine-like identity, lay along a transition whose direction recurred across cohorts, and mapped in situ to malignant NEPC territory. The relationships throughout are associative; the sections below keep one principal caveat at each claim and collect the remainder in the limitations.

Prior work framed the switch from adenocarcinoma to NEPC as lineage plasticity licensed by loss of *RB1* and *TP53* and supported by neural-lineage and pluripotency factors such as SOX2 [3,4]. Our data refine this picture by specifying the endpoint: the resulting state resembles a fetal rather than an adult neuroendocrine program. This suggests that the transition re-accesses a developmental transcriptional repertoire rather than assembling an arbitrary marker set, in line with phenotypic plasticity, disrupted differentiation, and oncofetal reprogramming as recurrent cancer themes [11,12]. A related logic has organized the SCLC field, where subtypes defined by ASCL1, NEUROD1, POU2F3, and YAP1 carry distinct regulatory programs [7,8]. The ASCL1-dominant assignment we observed aligns NEPC with aspects of ASCL1-high SCLC and points to a neuroendocrine regulatory logic shared across two epithelial cancers of different organs. Prior single-cell and chromatin studies have resolved ASCL1- and NEUROD1-defined NEPC subtypes and shown epigenetic convergence of NEPC with other neuroendocrine carcinomas and small-cell lung cancer [9,10,19]; our analysis adds a developmental dimension, assigning the ASCL1-defined state a fetal, lineage-restricted reference best supported as pulmonary neuroendocrine-like among the alternatives tested.

Among neuroendocrine lineages, the data favored a pulmonary neuroendocrine-like identity over islet-endocrine, chromaffin, and sympathoblast alternatives, with lung specifiers detectable and islet and sympathoblast specifiers near absent. An external specificity control reinforced this: a neuroblastoma reference — neuroendocrine and ASCL1-expressing but neural-crest in origin — mapped to chromaffin/neural-crest rather than pulmonary neuroendocrine identities under the same method (0/36, versus 29/30 for SCLC-A and 3/3 for NEPC), indicating the pulmonary-neuroendocrine assignment is not an artifact of mapping any neuroendocrine state to the lung lineage. The distinction between lung and islet was sensitive to method choice and associated with FOXA2: it held on regulons chosen to separate the two lineages and on most samples, whereas the two converged on the full shared regulon set. The most conservative reading is therefore a lineage-restricted identity within a neuroendocrine program with marker and regulatory features shared across organs, including pulmonary contexts linked to ASCL1 and pancreatic or other contexts linked to INSM1 [7,20], with pulmonary neuroendocrine as the best-supported member. These comparisons argue against treating the signal as generic neuronal or neuroendocrine dedifferentiation; it is more consistent with a constrained fetal neuroendocrine state, even if the boundary between pulmonary and islet remains provisional. One distinction matters for interpretation: “pulmonary neuroendocrine-like” denotes transcriptional resemblance, not an organ of origin, because NEPC arises in the prostate. Consistent with that, the reverse spatial mapping localized the program within fetal lung to the mature neuroendocrine compartment, a localization within the lung rather than evidence of pulmonary derivation.

The transition was consistent with an ordered sequence whose direction recurred across cohorts: factors associated with pioneer activity including FOXA2 and SOX2 reached their NEPC values early, while ASCL1, NEUROD1, and MYCN shifted late, with a monotonic trend in an independent staged series. This ordering is compatible with a model centered on pioneer factors in which FOXA family factors engage closed chromatin and prime regulatory elements ahead of lineage-defining factors [21]. In NEPC models, FOXA2 has also been reported to orchestrate the adeno-to-neuroendocrine transition and to be induced by androgen deprivation [16]. The ordering is cross-sectional, describing the arrangement of samples and pseudotime rather than elapsed time in any single tumor, so it should not be read as a timed developmental clock.

The regulatory data provide a candidate explanation for the pattern centered on ASCL1, consistent with recent work implicating ASCL1 in prostate cancer neuroendocrine differentiation [6], and resolve into three layers. At the expression layer, five orthogonal methods converged on a neuroendocrine circuit centered on ASCL1. At the chromatin layer, the convergent regulatory elements fell preferentially in fetal pulmonary neuroendocrine open chromatin rather than in non-neuroendocrine lung or a chromaffin decoy. At the factor layer, FOXA1, which was not differentially expressed and so invisible to the expression screen, occupied these elements more often than non-convergent ASCL1 peaks in both NEPC and adenocarcinoma models. With prior work showing that FOXA1 is redeployed from androgen receptor to neuroendocrine regulatory elements and is required for NEPC [15], these layers support a candidate model in which FOXA1 retained from the luminal state may provide a permissive chromatin context at fetal-neuroendocrine enhancers for a circuit centered on ASCL1. The model is nominated, not proven: the evidence is associative, rests on three NEPC patients, and draws its ASCL1 cistrome from SCLC/cell line models and its FOXA1 and H3K27ac profiles from LuCaP lines rather than primary patient NEPC.

These nominations are testable therapeutic hypotheses, not treatment recommendations. ASCL1 was the strongest selective dependency among the neuroendocrine regulators examined in a malignant neuroendocrine proxy, marking it as a candidate dependency while leaving open whether NEPC tumors share it. DLL3, a reported ASCL1 target in pulmonary neuroendocrine and small-cell lung cancer models [7], was upregulated here and is a reported, experimentally supported surface target in NEPC [17]; given the activity of DLL3-directed therapy in SCLC [22], it is the nominated node with the clearest existing translational rationale, though NEPC-specific clinical activity remains untested. The enhancer circuit involving FOXA1 and ASCL1 is conceptual rather than presently druggable and is offered as a direction for mechanistic work.

Several limitations bound these conclusions. The discovery NEPC sample is small (three patients), mitigated but not removed by cross-cohort replication of direction and effect size. inferCNV reports relative copy number without allele information, so the shared architecture is consistent with clonal relatedness without proving it. The forward spatial analysis rests on a single section from one patient and is exploratory, and the reverse mapping is at the region level and reflected shared neuroendocrine weighting rather than a weighting unique to NEPC. The cistrome and ChIP evidence comes from cell line and xenograft models, not primary patient NEPC, the principal gap between the nominated circuit and patient tissue.

Future work follows from these gaps. Key next steps include cistromes from primary patient NEPC and functional testing of FOXA1/ASCL1 dependence, including the enhancer circuit involving FOXA1 and ASCL1, in patient-derived models. Additional work should test whether the lung-islet ambiguity reflects a genuinely shared program or a reference limitation, and whether the in situ localization generalizes at cohort scale. In this set of cohorts, NEPC was best understood as a malignant, adenocarcinoma-related state that converges on a fetal, lineage-restricted, pulmonary neuroendocrine-like program, with an ordered transcriptional transition and a nominated regulatory architecture involving FOXA1 and ASCL1.

## Conclusions

NEPC is best understood as a malignant, adenocarcinoma-related state that converges on a fetal, lineage-restricted, pulmonary neuroendocrine-like program rather than a generic neuroendocrine one, reached through an ordered luminal-to-neuroendocrine transition and associated with a nominated regulatory circuit centered on ASCL1 with FOXA1 binding. Because these relationships are associative and rest on a small number of patients, with chromatin evidence from model systems rather than primary patient NEPC, functional testing in patient-derived models is the key next step. The study refines the NEPC neuroendocrine state from a generic phenotype to a specific developmental reference and nominates regulatory and surface candidates linked to ASCL1 for further study.

## Materials and Methods

### Data sources and cohorts

Effective n was defined as patients, samples, sections, or cell lines, not cells; cross-sample tests used per-patient pseudobulk aggregation. The discovery scRNA-seq cohort was GSE264573 (10x Genomics): 35,696 tumor cells from 20 patients (17 CRPC, 3 NEPC), including 24,089 adenocarcinoma and 11,607 NEPC cells. Per-patient analyses used the 13 patients with at least 100 tumor cells (10 adenocarcinoma, 3 NEPC). Replication and transition datasets were GSE210358 (three samples with NE features) and GSE137829 (six patients). Bulk cohorts were Beltran NEPC [1], SU2C/WCDT metastatic CRPC [23], TCGA prostate adenocarcinoma (TCGA-PRAD; [24]), and TARGET neuroblastoma [25]. Developmental references were Descartes fetal expression (GSE156793; [13]), Domcke fetal chromatin accessibility (GSE149683; [14]), He fetal lung expression/Visium (E-MTAB-11266; [18]), CZ CELLxGENE Census adult neuroendocrine references (release 2025-11-08; [26]), and Quach fetal lung [27]. Regulatory data included ASCL1 ChIP-seq in NEPC (GSE183200), SCLC-A (GSE69394), and neuroblastoma decoy cell lines (GSE120074); LuCaP FOXA1/H3K27ac ChIP-seq (GSE161948), ATAC-seq (GSE156291), and RNA-seq (GSE126078); and DepMap CRISPR dependencies (release 24q4). Spatial data included prostate NEPC Visium (GSE230282; one section), a NEPC foci Visium screen (GSE278936; 52 sections), He fetal lung Visium (seven sections), Quach fetal lung sections, and a mouse model Visium feasibility screen (GSE246762).

### Single-cell processing

GSE264573 was processed in Seurat [28]. Upstream quality control and normalization yielded 35,696 retained cells with nFeature_RNA 240–13,969, nCount_RNA 300–240,801, and mitochondrial content <25%. Integration used fastMNN [29] on 2,000 highly variable genes; the shared neighbor graph used the first 20 MNN dimensions (k = 30). Doublets were called per sample with scDblFinder [30] after excluding samples with <50 cells; all NEPC patients had doublet rates <10%. Datasets reprocessed in-house were log-normalized, except Xenium data (SCTransform). Anchor transfer and deep generative models used 2,000 highly variable genes.

### Copy-number and malignancy inference

Per-cell copy number was inferred with inferCNV [31] against an internal diploid immune/endothelial reference (n = 64 cells; hg38 GENCODE v27 gene order; cutoff 0.1; denoise on; hidden Markov model (HMM) off). Aneuploid cells were those with copy-number burden, defined as mean squared deviation from the reference mean profile, above the 95th reference percentile. Calls were cross-checked with CopyKAT [32], which concurred for the strongest-CNV group (NEPC-A) but under-called aneuploidy in adenocarcinoma and NEPC-N (Cohen’s κ ≈ 0; Fig. S1F), a known limitation with a small diploid reference; malignancy therefore rests on inferCNV and the shared chromosome-level copy-number architecture rather than on CopyKAT concordance. Because inferCNV lacks allele information, clonal relatedness was assessed by chromosome-level correlation of mean profiles.

### Fetal lineage mapping

NEPC cells were mapped to 77 Descartes fetal cell types in an open-world framework using three classifiers: Spearman centroid correlation (unassigned if the top-1 minus top-2 margin was <0.03), Seurat canonical correlation analysis (CCA) label transfer (dims 1:30; score <0.5 unassigned), and scVI/scANVI ([33,34]; 30 latent dimensions; maximum posterior <0.5 rejected). SingleR [35] was a secondary check. MetaNeighbor [36] was assessed but excluded because it misplaced control populations in this cross-platform task. Pseudobulk NEPC profiles were also correlated with fetal centroids, both pan-atlas and resolved by organ.

### Transcription factor and regulon activity

Transcription factor activity was inferred with decoupleR ULM [37] on CollecTRI [38]. Per-dataset adenocarcinoma baselines were subtracted, and P values came from 200 size-matched random regulons (two-sided permutation). De novo gene-regulatory networks used pySCENIC [39,40] with GRNBoost2 and cisTarget (hg38 v10 motifs) for the tumor-intrinsic regulons; an all-R GENIE3 [41] plus AUCell pipeline scored a parallel fetal reference SCENIC network. De novo subtype axes were retained only after held-out positive and shuffled-regulon negative controls.

### Per-patient pseudobulk differential expression

Raw counts were aggregated per patient with at least 100 cells and tested between NEPC (n = 3) and adenocarcinoma (n = 10) in DESeq2 [42] using design ∼ group (Wald test; prefilter at least 10 counts in at least 3 samples; FDR < 0.05). Log-fold changes were shrunk with apeglm [43]. Cross-cohort validation used rank-based scores rather than gene-level differential expression (see Lineage scoring).

### Lineage scoring and meta-analysis

Rank-based scores were used across normalizations: singscore [44] for bulk and UCell [45] for spatial and gene-set scoring. The fetal-minus-adult lung-neuroendocrine program was the fetal-minus-adult neuroendocrine z-score difference after subtracting a generic fetal-to-adult shift estimated from T cell controls. Lineage subtype scoring required at least 50 cells per patient. NEPC versus adenocarcinoma was tested by two-sided Mann–Whitney with Cliff’s delta; effects across three cohorts were pooled as Hedges’ g by random-effects meta-analysis (metafor, REML; [46]), reporting the pooled estimate, 95% CI, and I².

### Trajectory and ordering

Pseudotime for the single transdifferentiating tumor (GSE137829 patient P6) was inferred with Slingshot [47] from the most luminal cluster. TSCAN, tradeSeq, GeneSwitches, Palantir, and CellRank 2 [48–52] provided orthogonal trajectory and fate checks. Pseudotime was interpreted as ordering, not elapsed time. Morphological/transcriptional stages were defined as adenocarcinoma, early-NE, and mature-NE: in the discovery cohort GSE264573 supplied the adenocarcinoma and mature-NE samples and the partially transdifferentiated GSE137829 supplied the early-NE stage, and the cross-sectional axis that treated samples as stages pooled 21 samples (15 adenocarcinoma, 3 early-NE, 3 mature-NEPC). Ordered stage changes (adenocarcinoma < early-NE < mature-NE) were tested by Jonckheere–Terpstra (clinfun v1.1.5; [53]); the batch-robust composite that treated samples as stages used within-dataset percentile ranks.

### Cistrome and regulatory-element analysis

ASCL1 ChIP-seq reads were aligned to GRCh38 with Bowtie 2 [54], deduplicated, filtered for quality, with ENCODE blacklist regions removed, and peak called with MACS2 ([55]; q < 0.05).

Cell lines passing an ASCL1 E-box motif gate yielded a 5,531-peak ASCL1 cistrome; 1,210 peaks overlapped the SCLC-A consensus (shared NEPC/SCLC-A subset).

Overlap with Domcke fetal pulmonary neuroendocrine open chromatin [14] was tested against 100 width-matched random genomic shuffles (empirical z and one-sided permutation P). He fetal pulmonary neuroendocrine accessibility at these peaks was tested against width- and count-matched shuffled regions by one-sided Wilcoxon rank-sum test (P < 1 × 10⁻⁵⁰). Coordinates were converted between genome builds with rtracklayer/liftOver.

The 353 shared peaks within Domcke fetal pulmonary neuroendocrine chromatin were designated convergent enhancers and the remaining peaks non-convergent; convergent and non-convergent peaks were counted after lifting the ASCL1 cistrome to hg19 to match the deposited FOXA1 and H3K27ac ChIP (353 and 5,198 in the lifted coordinates, which therefore need not sum to the hg38 cistrome total). FOXA1 and H3K27ac occupancy at convergent versus non-convergent enhancers [15] was compared by paired Wilcoxon signed-rank test across NEPC LuCaP lines (FOXA1 87% vs 34%, P = 0.016; H3K27ac 92% vs 38%, P = 0.031). Beyond-ASCL1 motif enrichment used motifmatchr with JASPAR2022 [56,57]. Candidate regulators were also nominated by ChEA3 target enrichment [58], DepMap CRISPR dependency (Chronos gene effect, release 24q4; [59]), and ANANSE enhancer network influence [60] from LuCaP ATAC and RNA with GimmeMotifs [61].

### Spatial transcriptomics

Visium spots were deconvolved with robust cell type decomposition (RCTD) full mode [62] against discovery tumor cells plus immune/stromal references. Fetal lung cell state abundances used the published cell2location decomposition of the fetal lung atlas [18,63]. Region programs were scored with UCell, spatial transcription factor activity with decoupleR, and spatial autocorrelation with Moran’s I (k = 6 neighbors).

Malignant versus non-malignant discrimination used AUROC (pROC; [64]), before and after residualizing scores on per-spot complexity; significance used 200 label and weight permutations. Forward analysis tested colocalization of fetal pulmonary neuroendocrine and pan-neuroendocrine programs within the malignant region defined by copy number. Reverse analysis scored a neuroendocrine signature weighted by NEPC pseudobulk differential expression across fetal lung sections and ranked fetal neuroendocrine substates by complexity-controlled partial Spearman correlation with a 200-permutation weight permutation control. Xenium reverse mapping used Seurat anchor transfer.

### Statistics, software, and reproducibility

Significance was set at P or FDR < 0.05. Multiple testing used Benjamini–Hochberg, except the mapping test across 77 cell types (Bonferroni). Tests were two-sided unless a directional hypothesis was prespecified (mapping, specificity, and dependency controls). A fixed seed (42) was used for all stochastic steps; scANVI initialization sensitivity was additionally assessed across five seeds (1, 7, 13, 42, 99; Fig. S2G). Core transcription-factor and pseudobulk differential-expression findings (ASCL1 up, REST and AR down) were robust to leave-one-out exclusion of each NEPC patient; NEUROD1 remained directionally elevated but lost transcription-factor significance when HP21 was excluded (Fig. S1G). Analyses used R 4.5.3 (Bioconductor 3.22) and Python 3.11 for single-cell/spatial work; ANANSE used Python 3.10. Package versions queried on 2026-06-23 were: Seurat 5.4.0, scDblFinder 1.24.10, inferCNV 1.26.0, CopyKAT 1.1.0, decoupleR 2.16.0, AUCell 1.32.0, GENIE3 1.32.0, RcisTarget 1.29.0, SingleR 2.10.0, MetaNeighbor 1.28.0, DESeq2 1.50.2, singscore 1.30.0, UCell 2.14.0, metafor 5.0.1, pROC 1.19.0.1, slingshot 2.18.0, TSCAN 1.48.0, tradeSeq 1.24.0, GeneSwitches 1.0.0, clinfun 1.1.5, spacexr/RCTD 2.2.1, motifmatchr 1.32.0, TFBSTools 1.48.0, JASPAR2022 0.99.8; scvi-tools 1.4.2, scanpy 1.11.5, anndata 0.12.16, Palantir 1.4.4, CellRank 2.0.7, cellxgene-census 1.17.0; Bowtie 2 2.5.5, MACS2 2.2.9.1, SAMtools 1.23.1, BEDTools 2.31.1, deepTools 3.5.6, ANANSE 0.5.1, GimmeMotifs 0.18.3, apeglm 1.32.0, and pySCENIC 0.12.1 (Python 3.10).

## Data and code availability

All datasets were previously published, public, and de-identified; no new human or animal data were generated, and no additional ethical approval was required. Primary sources were GEO (GSE264573, GSE210358, GSE137829, GSE183200, GSE69394, GSE120074, GSE156793, GSE149683, GSE161948, GSE156291, GSE126078, GSE230282, GSE278936, GSE246762), ArrayExpress (E-MTAB-11266), TARGET neuroblastoma, DepMap release 24q4, and CZ CELLxGENE Census release 2025-11-08. Analysis code, consisting of custom R and Python scripts around the published tools above, will be deposited in a public repository and archived with a persistent digital object identifier (DOI) upon publication, and is available from the corresponding author on reasonable request during review.

## Ethics approval and consent to participate

This study analyzed publicly available, de-identified datasets only; no new human or animal data were generated and no additional ethical approval was required. Consent for publication is not applicable.

## CRediT authorship contribution statement

**Wenchang Yue:** Conceptualization, Methodology, Software, Formal analysis, Investigation, Data curation, Visualization, Writing original draft, Writing review and editing. **Natasha Kyprianou:** Writing review and editing. **Ashutosh K. Tewari:** Funding acquisition. **Babu J. Padanilam:** Conceptualization, Supervision, Funding acquisition, Project administration, Resources, Writing review and editing.

## Data availability

All datasets analyzed in this study are previously published and publicly available; accession numbers are listed under Materials and Methods (Data and code availability). The custom R and Python analysis code will be deposited in a public repository and archived with a persistent DOI upon publication, and is available from the corresponding author on reasonable request during review.

## Acknowledgements

The authors thank The Cancer Genome Atlas (TCGA) Research Network, the Stand Up to Cancer–Prostate Cancer Foundation (SU2C-PCF) Dream Team, the developmental and human cell atlas consortia, and all contributors to the Gene Expression Omnibus (GEO), ArrayExpress, CZ CELLxGENE, and DepMap for making their data publicly available.

