## Supplementary Information for "Neuroendocrine prostate cancer converges on a fetal pulmonary neuroendocrine-like program"

This file contains Supplementary Figures S1–S6 and their legends. Supplementary figures are referenced in the main text.

### **Supplementary figures**

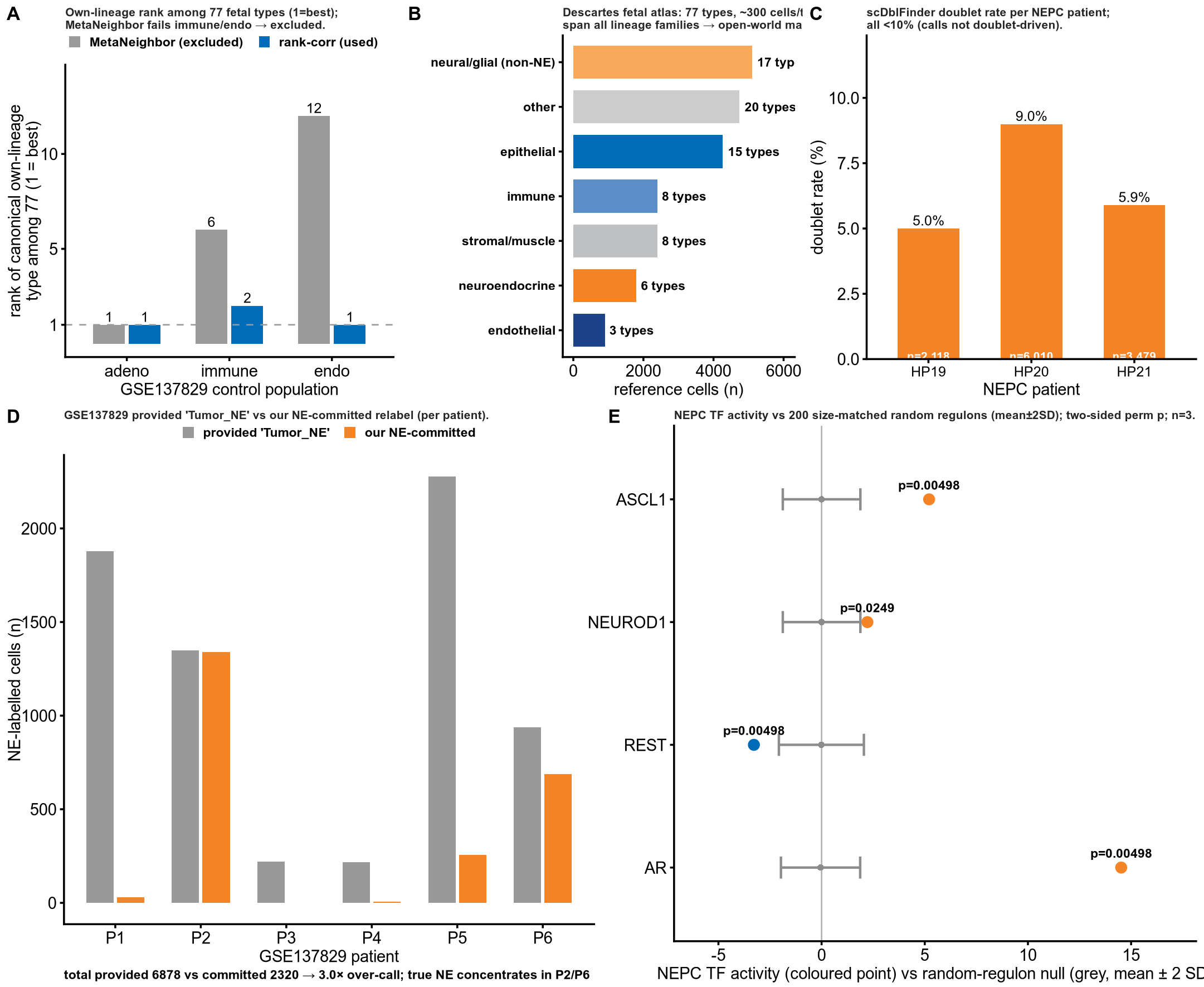

**Figure S1. Identity-mapping rigor and single-cell QC.** **(A) Why MetaNeighbor is excluded.** Own-lineage rank of each control among the 77 fetal types (1 = best): rank-correlation places immune at Lymphoid and endothelium at Vascular-endothelial (rank 1–2), whereas MetaNeighbor fails them (rank 6/77 and 12/77) and is not trusted on this cross-platform task. n = GSE264573 control populations. **(B) Reference composition.** The full Descartes atlas (77 cell types, approximately 300 cells/type) spans all families, so NEPC is mapped open-world, not into a neural-only reference. **(C) Doublet QC.** scDblFinder rate per NEPC patient all < 10%. n = 3 NEPC patients (HP19/20/21). **(D) GSE137829 over-call.** The provided Tumor_NE label over-calls NE approximately 3× vs our committed relabeling (6,878 to 2,320 cells), concentrated in P2/P6, indicating an early/partial transition state, not mature NEPC. n = 6 GSE137829 patients. **(E) TF-activity permutation control.** Observed NEPC activity of ASCL1, NEUROD1, REST, AR vs 200 size-matched random regulons (mean ± 2 SD); all outside the null (|z| 2.35–15.2; perm p = 0.005 / 0.025 / 0.005 / 0.005), indicating a regulon-specific readout. n = 3 NEPC patients. **(F) inferCNV vs CopyKAT malignancy concordance.** Per-group fraction of cells called aneuploid by inferCNV vs CopyKAT (n = 4,960 cells); the methods concur on NEPC-A but CopyKAT under-calls adenocarcinoma and NEPC-N (Cohen's κ ≈ 0), so malignancy rests on inferCNV plus the shared chromosome-level copy-number architecture, not CopyKAT concordance. **(G) Leave-one-out sensitivity.** Core findings re-tested dropping each NEPC patient (n = 2 vs 10 adenocarcinoma per fold); red = full cohort, grey = the three folds; filled = P < 0.05, open = ns. ASCL1 (up), REST and AR (down) hold sign and significance in every fold; NEUROD1 stays directionally up but its TF-activity significance is HP21-sensitive (perm P 0.025→0.065). n = 3 NEPC, 10 adenocarcinoma.

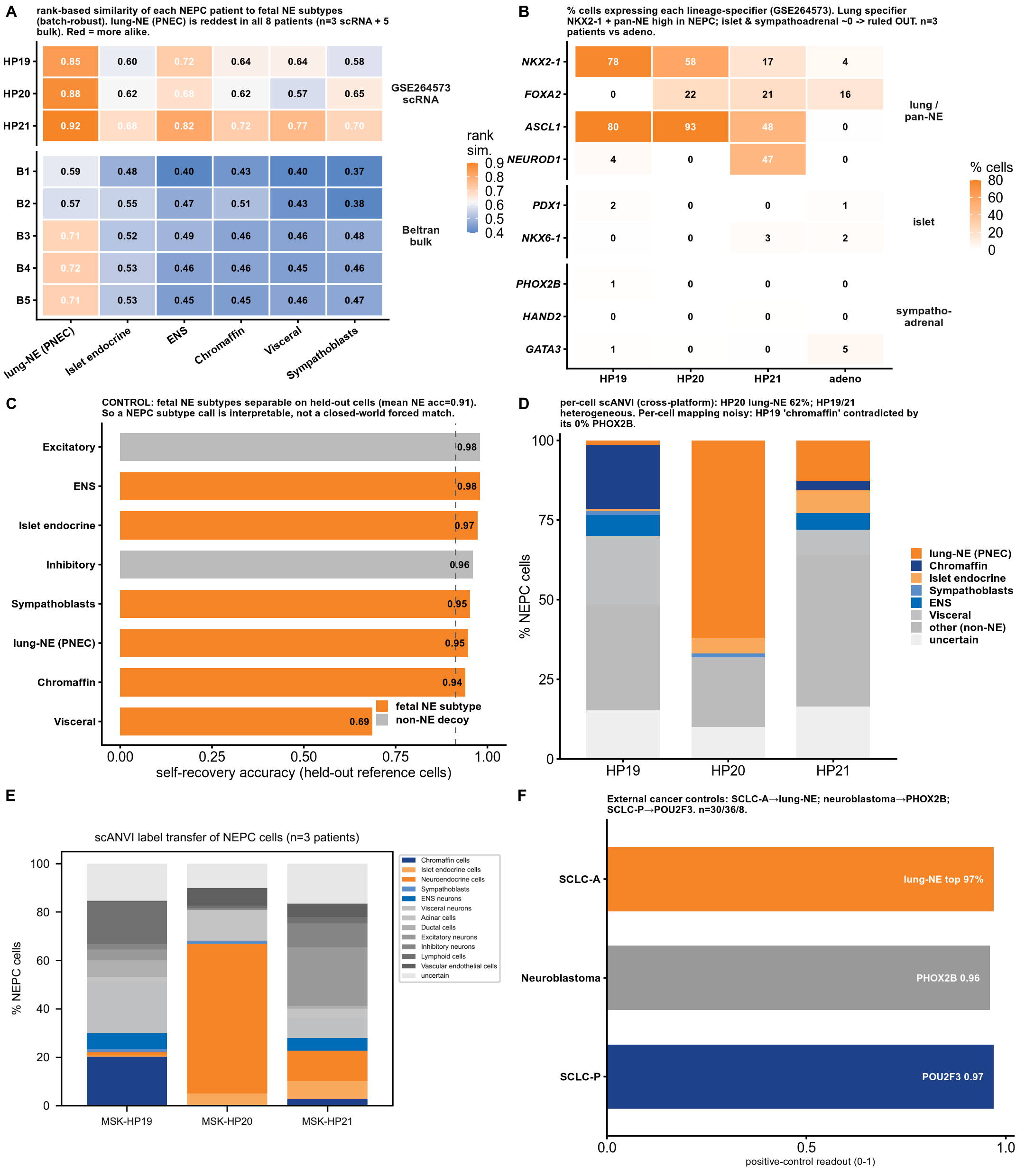

**Figure S2. Fetal-NE subtype resolution with external positive controls.** Panel D uses the earlier scANVI run and Fig 3D the later one, so fractions differ (method sensitivity, not duplicates). **(A) Subtype mapping.** Rank-based similarity to the 6 fetal NE subtypes: lung-NE/PNEC is the top match in all 8 samples (3 scRNA patients + 5 Beltran bulk), above adeno. **(B) Lineage specifiers (% cells).** Lung NKX2-1/TTF1 in 16 to 78% of NEPC vs 4% adeno; islet PDX1/NKX6-1 approximately 0 to 2.6%; sympathoblast PHOX2B/HAND2/GATA3 approximately 0%, ruling out islet and neural-crest/sympathoblast. **(C) Discriminability control.** Held-out fetal NE cells re-recovered at approximately 0.91 accuracy (clean islet/lung-NE/chromaffin confusion). n = 6 fetal NE subtypes. **(D) scANVI subtype fractions (per-cell).** Earlier run: HP20 majority lung-NE (61.9%), chromaffin approximately 0%; HP19/HP21 heterogeneous. n = 3 patients. **(E) scANVI raw (per-cell).** Per-cell assignments behind (D). n = 3 patients. **(F) External positive and specificity controls.** Under the same fetal-subtype mapper (discriminability gate passed), SCLC-A mapped to fetal pulmonary neuroendocrine in 29/30 lines (97%) and NEPC in 3/3 patients, whereas neuroblastoma — a neuroendocrine, ASCL1-expressing tumor of neural-crest origin — mapped to chromaffin and other neural-crest neuroendocrine lineages (0/36 to pulmonary neuroendocrine; n = 36 TARGET-NBL; PHOX2B = 0.96); SCLC-P POU2F3 = 0.97 (n = 8 cell lines). The pulmonary-neuroendocrine assignment was therefore lineage-specific, not a default for any neuroendocrine or ASCL1-driven state. **(G) scANVI vs random seed.** Per-patient scANVI fraction under five seeds (1, 7, 13, 42, 99); confident neuroendocrine-family (left) and fetal lung-neuroendocrine (right). Each point is one seed; bar = median; seed 42 (Fig. 3D) in red. HP20 is stable (median lung-neuroendocrine 95%), HP21 intermediate (35%), and HP19 seed-unstable (lung-neuroendocrine 1–51%; confident neuroendocrine 10–68%), so the per-patient identity call rests on the deterministic rank-correlation and anchor-transfer mappings (Fig. 3C, 3E). n = 3 NEPC patients (HP19–21), 5 seeds.

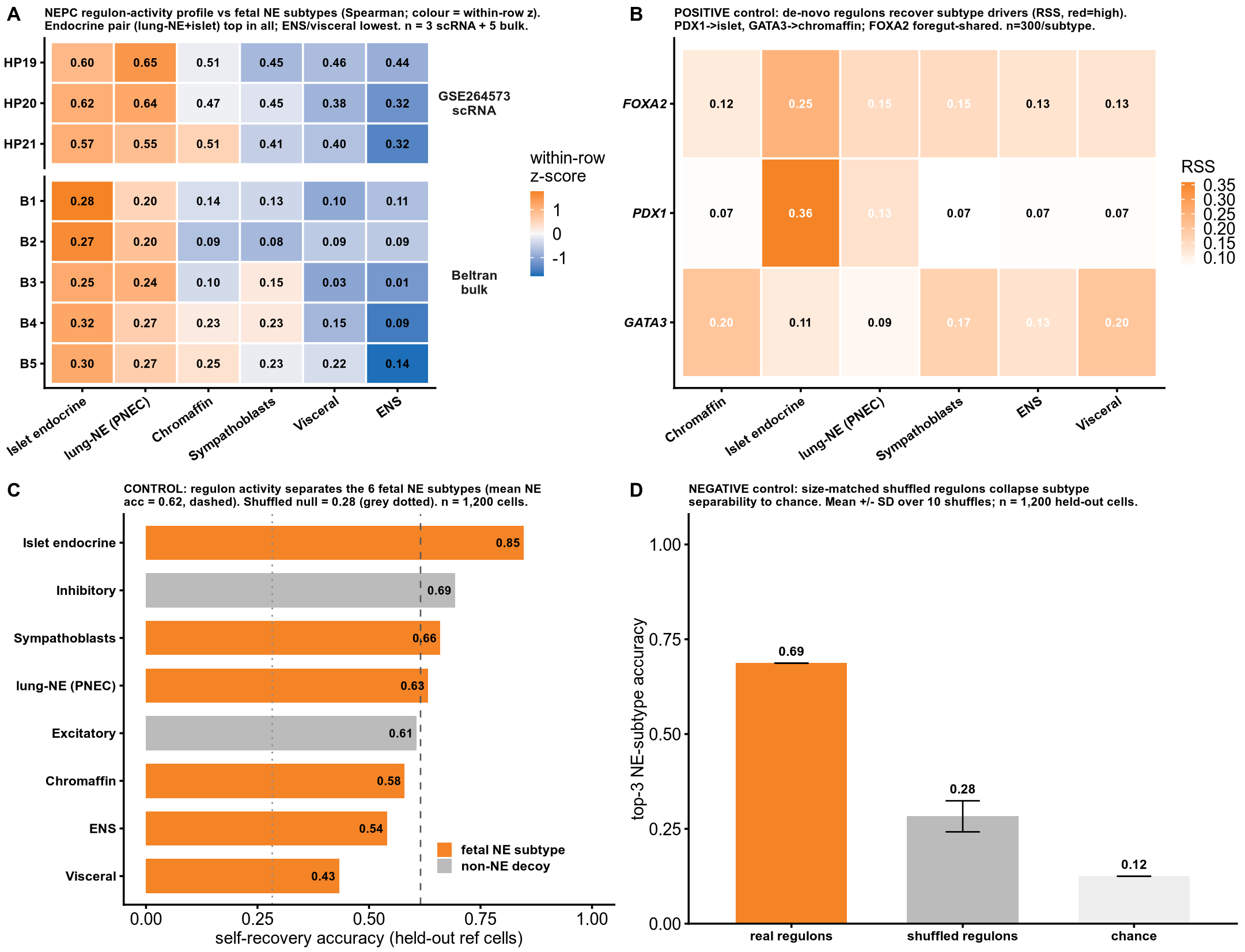

**Figure S3. Regulon-activity functional axis with controls.** Orthogonal de-novo regulon-activity line (weak–moderate independent support). **(A) Regulon mapping.** NEPC de-novo regulon-activity profile vs the 6 fetal NE subtypes: the endocrine pair (lung-NE + islet) is top-2 in all 8 samples; enteric/visceral and neural-crest excluded. n = 3 GSE264573 patients + 5 Beltran samples. **(B) Positive control (regulon specificity score [RSS]).** Recovers known drivers: PDX1 top-specific for islet (RSS 0.36); GATA3 tied top for chromaffin/visceral (0.203 / 0.203); FOXA2 foregut-shared. n = 300 cells/subtype. **(C) Discriminability control.** Per-subtype self-recovery accuracy on held-out cells (n approximately 1,200 cells). **(D) Negative control.** Shuffled regulons collapse separability (real top-3 accuracy 0.69 vs shuffled 0.28 ± 0.04), indicating regulon specificity.

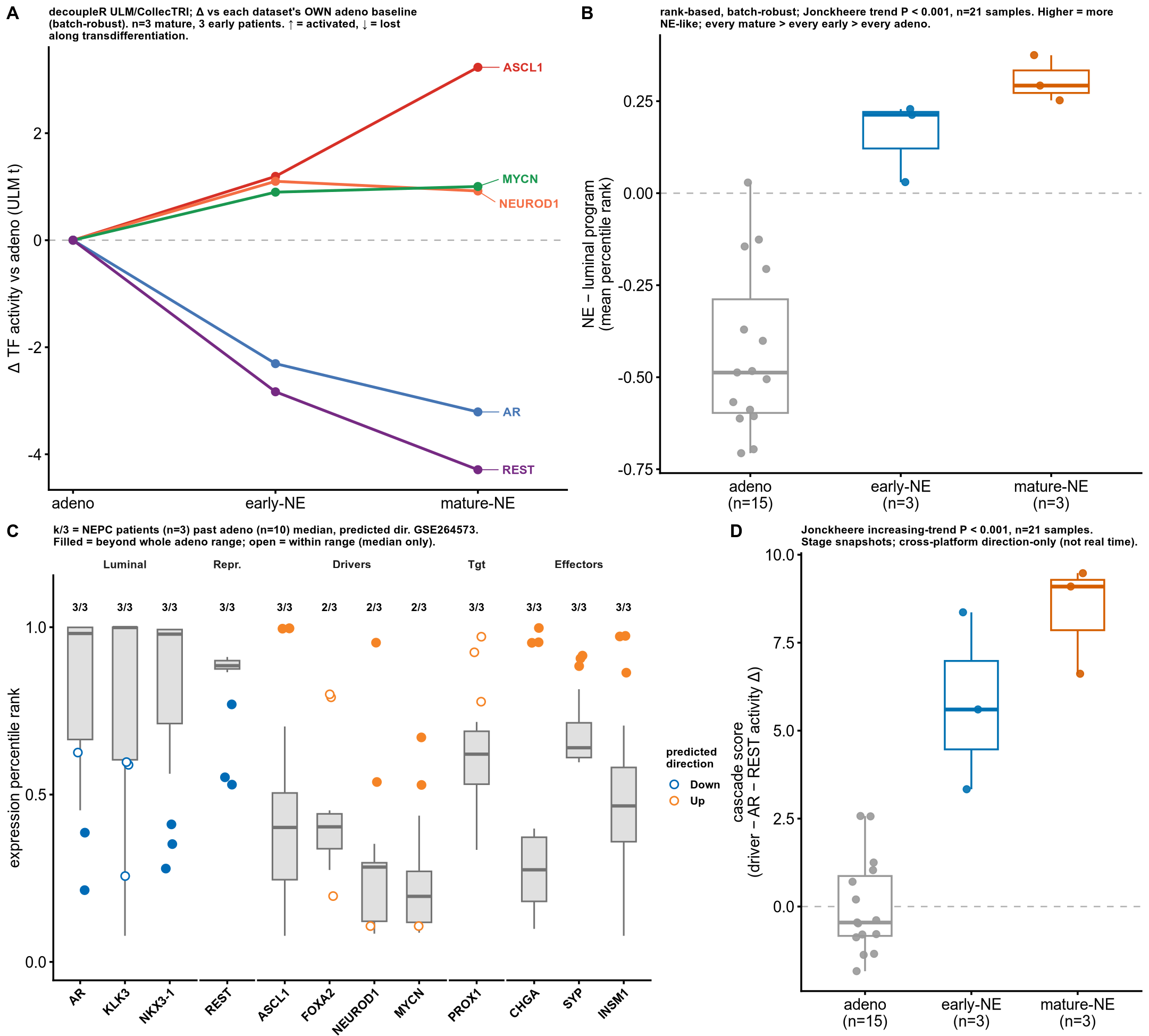

**Figure S4. Transdifferentiation trajectory and cascade detail.** Relative ordering (Jonckheere), not real time; effective n = patients. **(A) Staging TF activity.** decoupleR ULM/CollecTRI Δ across the adeno, early-NE and mature-NE stages (each dataset vs its own adeno): ASCL1 +3.23, NEUROD1 +0.92, MYCN +1.0 rise; AR −3.21 (control) and REST −4.29 fall. n = 3 mature + 3 early + adeno patients. **(B) Staging marker balance.** Rank-based NE−luminal balance; Jonckheere increasing-trend P < 0.001 (n = 21 samples); every mature > every early > every adeno. **(C) Per-patient cascade rank.** Strong nodes move past the adeno range in 3/3 NEPC patients. **(D) Ordered cascade score.** Composite adeno < early-NE < mature-NE (0, 5.77, 8.40; Jonckheere P < 0.001, n = 21 samples; robust to dropping FOXA2).

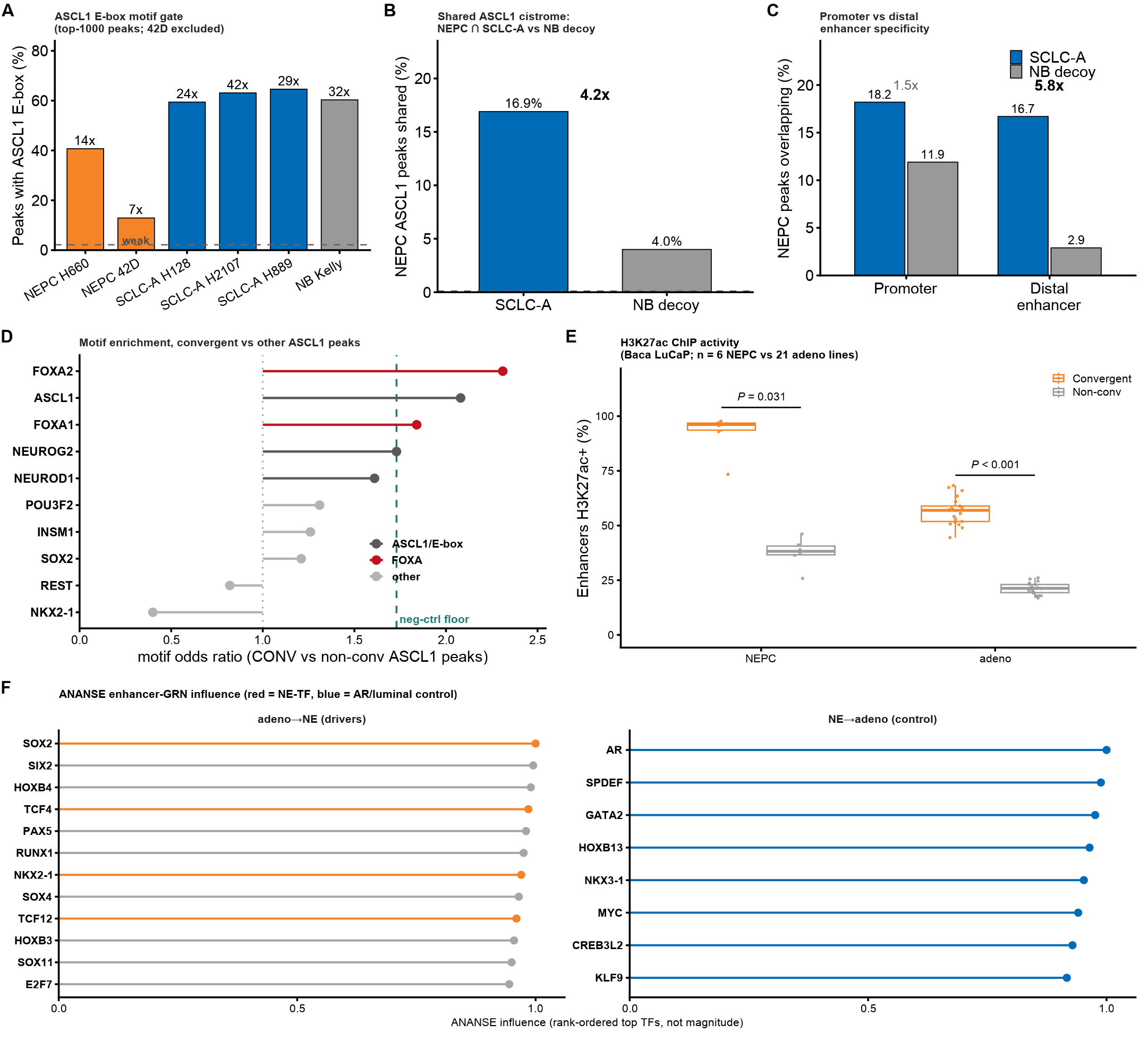

**Figure S5. ASCL1 cistrome QC and candidate-regulator cross-validation.** Cistrome data are cell-line/PDX and region-level, with no patient evidence; the 3-patient cap is unchanged. **(A) ASCL1 E-box motif gate.** Top-1000 ASCL1 peaks with the E-box (JASPAR MA1100.2) vs matched-random: NEPC-H660 41% (14×, pass); 42D fails (13%, 6.8×) and is excluded, so the NEPC cistrome rests on H660 alone. **(B) Lineage specificity.** Shared NEPC/SCLC-A 16.9% vs shared NEPC/neuroblastoma-decoy 4.0% = 4.2×, well above shuffled null 0.1%. **(C) Promoter vs distal.** Enhancer-concentrated: distal SCLC 16.7% vs decoy 2.9% = 5.8× vs promoters 1.5×. **(D) FOXA co-motif.** In convergent vs non-convergent ASCL1 peaks, FOXA is the only robust beyond-ASCL1 motif: FOXA2 OR = 2.31, FOXA1 OR = 1.84, ASCL1 E-box OR = 2.08, above the negative-control floor 1.73. Nominates a FOXA pioneer co-factor (validated as FOXA1 binding in Fig. 5E). n = 353 convergent peak regions (cell-line models, not patient tissue). Motif is not binding. **(E) H3K27ac.** Convergent enhancers active: 92.2% vs 37.6% in NEPC (6 lines, paired P = 0.031); 56.7% vs 21.2% in adeno (21 lines, P < 0.001). Baca GSE161948 LuCaP. **(F) ANANSE enhancer-GRN.** The NE-to-adeno control returns AR (rank 1), SPDEF and GATA2; adeno-to-NE returns SOX2 (rank 1), TCF4 (rank 4) and NKX2-1 (rank 7) (ASCL1/NEUROD1 via TCF4/TCF12 = E-box limitation). Rank, not magnitude; cell-line/PDX.

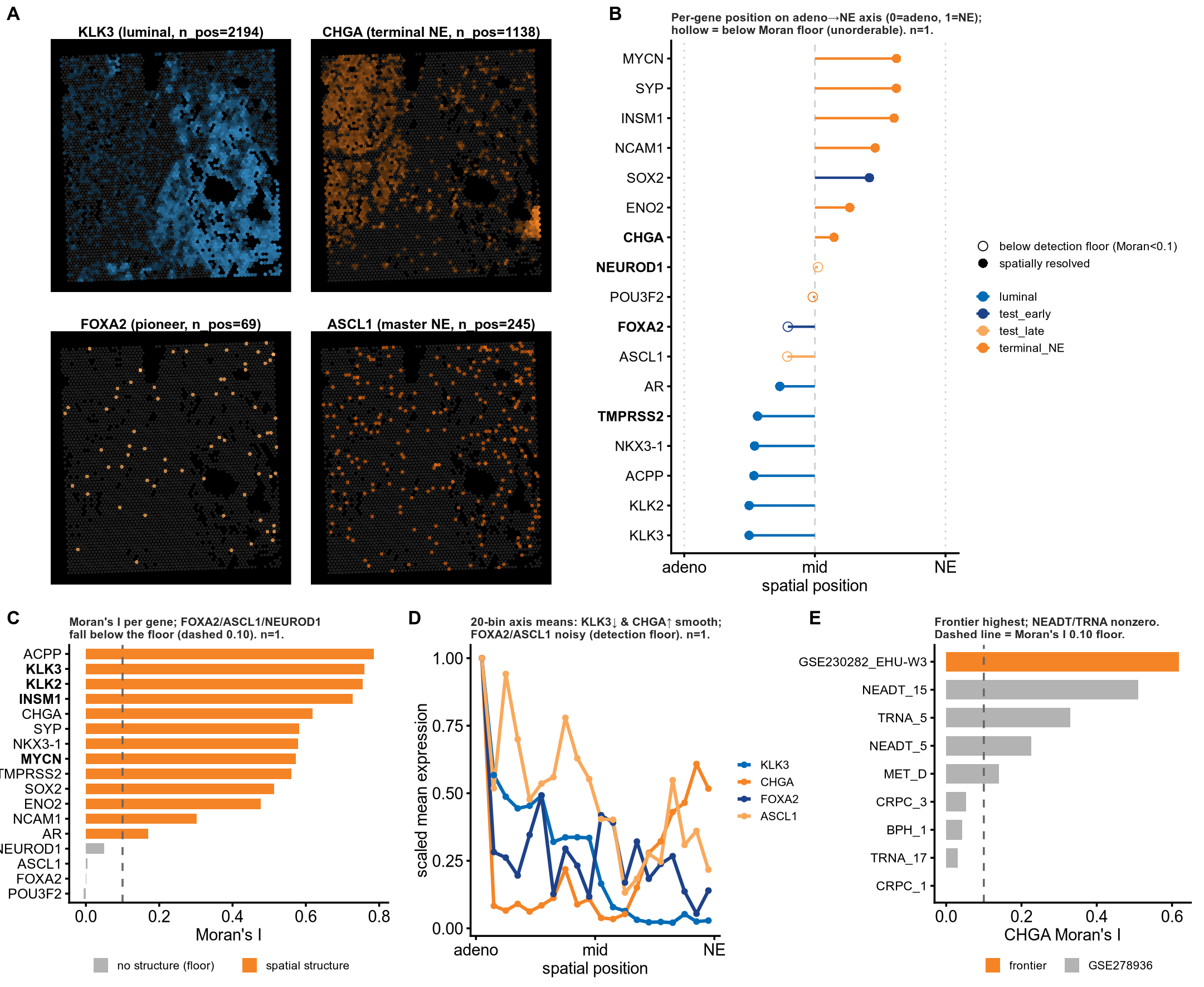

**Figure S6. In situ adenocarcinoma-to-neuroendocrine spatial axis (exploratory; n = 1 section, 1 patient).** In this NEPC Visium section (4,397 spots), the adenocarcinoma-to-neuroendocrine axis is spatially continuous, but lineage TFs fall below the Visium detection floor and are not spatially resolvable. **(A) Gene-domain maps.** KLK3 (luminal, n_pos 2,194) and CHGA (NE, n_pos 1,138) form complementary domains; FOXA2 (69 spots) and ASCL1 (245 spots) are sparse. **(B) Per-gene axis position.** 0 = adeno, 1 = NE; hollow = below the Moran floor (FOXA2/ASCL1/NEUROD1). **(C) Moran’s I per gene.** FOXA2/ASCL1/NEUROD1 fall below the 0.10 floor (0.001 / 0.005 / 0.05), so their spatial position is noise. **(D) 20-bin axis profiles.** KLK3 decreases and CHGA increases smoothly; FOXA2/ASCL1 noisy. Resolvable markers structured (KLK3 Moran’s I = 0.759; CHGA = 0.619); the luminal and NE programs were anti-correlated (ρ = −0.38). **(E) Negative control.** CHGA structure highest at the real frontier (Moran’s I = 0.619); many non-frontier sections low (BPH_1 = 0.042, CRPC_3 = 0.053) but not all zero (NEADT_15 = 0.510, TRNA_5 = 0.329), i.e. a specificity screen, not proof of absence.
